# FoxP3 can fold into two distinct dimerization states with divergent functional implications for T cell homeostasis

**DOI:** 10.1101/2022.02.08.479534

**Authors:** Fangwei Leng, Wenxiang Zhang, Ricardo N. Ramirez, Juliette Leon, Yi Zhong, Joris van der Veeken, Alexander Y. Rudensky, Christophe Benoist, Sun Hur

**Author notes:** These authors contributed equally (Fangwei Leng, Wenxiang Zhang).

## Abstract

FoxP3 is an essential transcription factor (TF) for immunologic homeostasis, but how it utilizes the common forkhead DNA-binding domain (DBD) to perform its unique function remains poorly understood. We here demonstrate that, unlike other known forkhead TFs, FoxP3 forms a head-to-head dimer using a unique linker (Runx1-binding region, RBR) preceding the forkhead domain. Head-to-head dimerization confers distinct DNA-binding specificity and creates a docking site for the cofactor Runx1. RBR is also important for proper folding of the forkhead domain, as truncation of RBR induces domain-swap dimerization of forkhead, which was previously considered the physiological form of FoxP3. Rather, swap-dimerization impairs FoxP3 function, as demonstrated with the disease-causing mutation R337Q, while a swap-suppressive mutation largely rescues R337Q-mediated functional impairment. Altogether, our findings suggest that FoxP3 can fold into two distinct dimerization states: head-to-head dimerization representing functional specialization of an ancient DBD and swap-dimerization with impaired functions.

## Introduction

The human genome is estimated to encode approximately 1,600 transcription factors (TFs). The vast majority of these TFs, however, utilize one of just 10 types of DNA binding domains (DBDs) (Lambert et al., 2018), many of which display remarkably conserved and narrow sequence specificity (Nitta et al., 2015). The apparent simplicity in DBD composition is in sharp contrast to the complex network of genes they control. How DBDs can divergently evolve to carry out distinct functions remains incompletely understood (Badis et al., 2009; Jolma et al., 2015; Reiter et al., 2017).

One of the largest families of DBDs in eukaryotes is the forkhead DBD, which commonly displays a winged-helix fold and recognizes the consensus sequence known as forkhead motif (FKHM) – TGTTTAC (Dai et al., 2021).There are about 50 forkhead TFs in human, which play important roles in key biological processes, from development to reproduction, to aging and to immunity (Benayoun et al., 2011; Hannenhalli and Kaestner, 2009). Some forkhead TFs function as pioneering TFs that can directly recognize DNA sequence within the condensed chromatin and open its structure with the help of chromatin modifiers (Drouin, 2014), while others utilize pre-existing chromatin landscape without dramatically altering its structure (Samstein et al., 2012). This suggests that there is great functional diversity even among the forkhead TFs not only in biological processes they control, but also in molecular mechanisms they employ.

FoxP3 is a forkhead TF that plays a critical role in development of regulatory T (Treg) cells, a branch of CD4^+^ T cells that suppress a variety of immune functions to prevent autoimmunity and excessive inflammation (Bennett et al., 2001; Brunkow et al., 2001; Fontenot et al., 2003; Hori et al., 2003). Earlier studies showed that certain mutations in FoxP3 lead to the multiorgan autoimmune disease immune dysregulation, polyendocrinopathy, enteropathy, X-linked (IPEX) syndrome in human and similar autoimmune conditions in mouse (Bennett et al., 2001; Brunkow et al., 2001; Chatila et al., 2000; Gambineri et al., 2008; Rubio-Cabezas et al., 2009; Tanaka et al., 2005; Wildin et al., 2001). FoxP3 is key to determining and maintaining the Treg identity by transcriptionally up- or down-regulating hundreds of genes (Kwon et al., 2017; van der Veeken et al., 2020). Despite its importance in immune homeostasis, molecular functions of FoxP3 remain poorly understood. For examples, it is highly debated whether FoxP3 is a transcriptional activator or suppressor, or can be both depending on the target genes (Zheng et al., Nature, 2007; Arvey et al., 2014; Kwon et al., 2017; Li et al., 2007; Zemmour et al., 2021); and which of the genes affected by FoxP3 are the direct target genes versus those regulated indirectly, perhaps mediated by FoxP3’s direct target genes (Ricardo N. Ramirez, In Press; van der Veeken et al., 2020; Zemmour et al., 2021). It is also unclear how FoxP3 alters the target gene expression as FoxP3 binds predominantly to genomic loci that have pre-established chromatin accessibility and induces little change in the accessibility of the bound sites (Samstein et al., 2012; van der Veeken et al., 2020; Yoshida et al., 2019).

Equally puzzling are the biochemical and structural properties of FoxP3. FoxP3 contains an N-terminal proline-rich region that recruits a variety of cofactors, zinc finger (ZF) and coiled coil (CC) that forms an antiparallel dimer (Song et al., 2012) (Figure 1A). Followed by CC is a long linker that recruits the cofactor Runx1 (Ono et al., 2007)(to be named Runx1-binding region; RBR) and the forkhead domain that is responsible for DNA binding. Among these domains, the forkhead domain is most frequently mutated in IPEX patients and has been most extensively studied (Barzaghi et al., 2012; Huang et al., 2020). Previous studies reported that isolated FoxP3 forkhead folds into an unusual domain-swap dimer that drastically differs from the winged-helix monomeric structure typical of forkhead DBD (Bandukwala et al., 2011; Chen et al., 2015).

**Figure 1.**
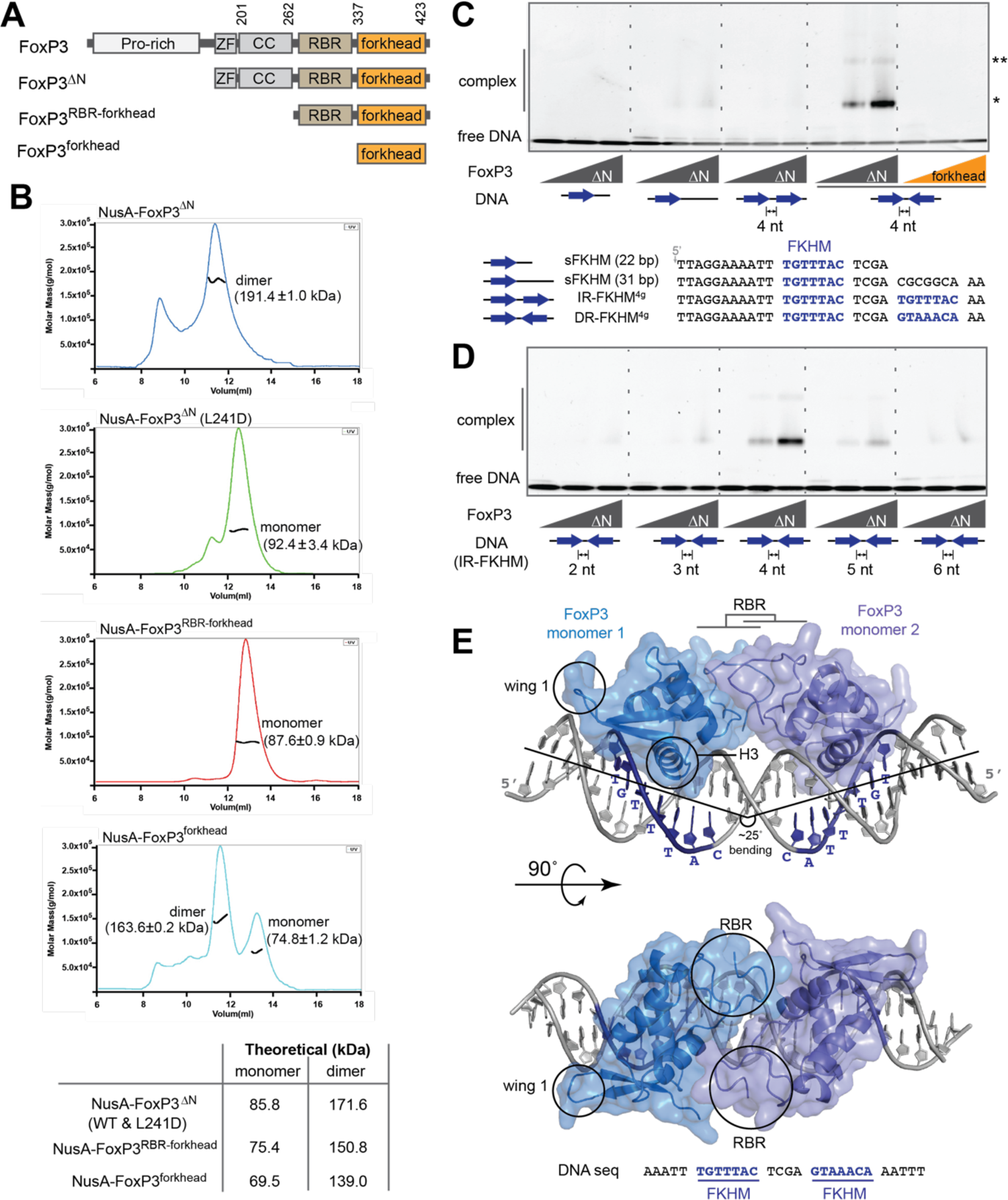
Overall architecture of FoxP3. See also Figures S1 & S2. A. Domain architecture of FoxP3^ΔN^, FoxP3^RBR-forkhead^ and FoxP3^forkhead^. (ZF: zinc finger; CC: coiled coil; RBR: Runx1-binding region). B. SEC-MALS of various truncation variants of FoxP3. Experimentally determined mass values are shown in parenthesis on the graphs. Theoretical values are shown in the table below. The NusA tag was fused for all constructs to increase the accuracy of mass estimation. L241D disrupts CC dimerization. C. EMSA of FoxP3^ΔN^ or FoxP3^forkhead^ (0, 0.4 and 0.8 μM) using four DNA oligos (0.2 μM) with different FKHM arrangements. Sybr Gold stain was used to visualize DNA. While FoxP3^ΔN^ binds DNA predominantly as a dimer (*), a small population of higher-order oligomer (**) was also seen. D. EMSA of FoxP3^ΔN^ (0, 0.4 and 0.8 μM) with DNA containing IR-FKHM (0.2 μM) with varying gap sizes. E. Crystal structure of Foxp3^ΔN^ in complex with IR-FKHM^4g^ DNA. Two non-swap FoxP3 monomers form a head-to-head dimer through the RBR loop. ZF and CC were present in the crystal (See Figure S2E), but were not resolved. The helix 3 (H3) and wing 1 that are characteristics of the canonical forkhead structure are indicated with circles. Data in (B-D) are representative of at least three independent experiments.

Closely related TFs, FoxP1 and FoxP2, were also found to form domain-swap dimers (Chu et al., 2011; Stroud et al., 2006), which led to the widely accepted notion that FoxP TFs may have evolved to adopt the swap-dimeric fold. However, others reported that a peptide that binds FoxP3 ZF-CC can disrupt FoxP3 dimerization, inconsistent with the domain-swap model of forkhead (Lozano et al., 2017). FoxP2 was also crystallized as a non-swap monomer as well as the domain-swap dimer (Stroud et al., 2006; Wu et al., 2006), further raising the question about the physiologically relevant structure of the FoxP forkhead domains. More globally, the overall architecture of FoxP3 beyond its DBD and how FoxP3 differs from other closely related TFs, such FoxP1 that also expresses in Tregs (Ghosh et al., 2018; Konopacki et al., 2019), to carry out its unique function also remains unclear.

Towards our goal of defining the overall architecture of FoxP3, we employed a combination of X-ray crystallography, biochemistry and functional assays in cells and mouse models. Our results suggest that FoxP3 forkhead, as well as those of other FoxP TFs, folds into the winged-helix, non-swap conformation, and that this requires the presence of the RBR linker. Individually folded forkhead domains then form a head-to-head dimer upon DNA binding. While other FoxP TFs also utilize their RBR-like linkers for monomeric folding, head-to-head dimerization is unique to FoxP3 and imparts distinct functions to FoxP3. Our functional data further provide new insights into the pathogenic mechanism for IPEX mutations and a new framework of understanding for FoxP3 functions.

## Results

### Overall architecture of FoxP3

We first attempted purifying full-length mouse FoxP3 protein from *E. coli*, but found it heavily degraded in the N-terminal Pro-rich region. This is consistent with multiple structural predictions (e.g. AlphaFold2, Jpred4) suggesting that the N-terminal region is intrinsically disordered (Drozdetskiy et al., 2015; Jumper et al., 2021). We therefore purified an N-terminal truncation variant (FoxP3^ΔN^) harboring ZF, CC, RBR and forkhead domains (Figures 1A and S1A). We also purified FoxP3^RBR-forkhead^ and FoxP3^forkhead^ for comparison. To confirm previous reports that FoxP3^forkhead^ constitutively forms a swap dimer (Bandukwala et al., 2011), we examined the oligomeric states of the purified FoxP3 proteins using size exclusion chromatography-coupled multiangle light scattering (SEC-MALS). We used FoxP3 fused with the protein tag NusA (60 kDa) to improve accuracy of molecular weight estimation by SEC-MALS. The result showed that FoxP3^forkhead^ was largely dimeric, consistent with the previous report (Figure 1B). To our surprise, FoxP3^RBR-forkhead^ was monomeric, which is incompatible with the swap dimeric structure of forkhead (Figure 1B). FoxP3^ΔN^ was a dimer but disruption of the CC dimerization by a mutation in the CC dimeric interface (L241D) (Song et al., 2012) converted it to a monomer (Figure 1B), suggesting that FoxP3 dimerization is exclusively mediated by CC, and that forkhead within the FoxP3^ΔN^ construct is a monomer.

Given that the observed differences in the oligomeric state of FoxP3 forkhead depends on other domains, we postulated that there may be different conformations of forkhead. We next asked how these different conformations affect DNA binding. We first examined DNA containing only a single FKHM (sFKHM, 22 or 31 bp) by electrophoretic mobility shift assay (EMSA), but found that binding was nearly undetectable for all three proteins of FoxP3^ΔN^, FoxP3^RBR-forkhead^ and FoxP3^forkhead^ up to 0.8 μM (Figures 1C, S1D and S1E). A previous study reported that a tandem repeat of FKHMs increased FoxP3 affinity (Koh et al., 2009). This prompted us to examine the potential effect of tandem repeats of FKHM, both direct repeats (DR) and inverted repeats (IR), with various gap sizes. The results showed that FoxP3^ΔN^ had markedly strong preference for IR-FKHM with 4 nt gap (IR-FKHM^4g^) (Figures 1C and 1D). We did not observe high affinity binding to DR-FKHM regardless of the gap size (Figures S1B). The same DNA specificity was observed for full-length FoxP3 ectopically expressed in 293T cells, suggesting that FoxP3’s preference for IR-FKHM^4g^ is independent of the source of the protein (Figure S1C).

Intriguingly, FoxP3^RBR-forkhead^ also displayed the same preference for IR-FKHM^4g^ (Figure S1D), while FoxP3^forkhead^ did not (Figure S1E). Note that FoxP3^forkhead^ was significantly less efficient in DNA binding than FoxP3^ΔN^ and FoxP3^RBR-forkhead^ (Figure 1C, last 6 lanes, and Figure S1D-E). Considering that the preference for IR-FKHM^4g^ was observed only with FoxP3^ΔN^ and FoxP3^RBR-forkhead^, which harbored monomeric forkhead, but not with swap-dimeric FoxP3^forkhead^, the different DNA selectivity is likely a direct consequence of different forkhead conformations. In fact, modeling suggests that the two swap dimers cannot simultaneously occupy two FKHMs in IR-FKHM^4g^ (Figure S1F), explaining why FoxP3^forkhead^ does not display the same preference for IR-FKHM^4g^ as with FoxP3^ΔN^ and FoxP3^RBR-forkhead^. These results strongly indicate that FoxP3 forkhead in more native domain architecture (FoxP3^ΔN^ and FoxP3^RBR-forkhead^) is not in the swap dimeric conformation, and that the true conformation of FoxP3 is the key to understanding its DNA specificity.

We thus co-crystallized FoxP3^ΔN^ in complex with IR-FKHM^4g^. For crystallization, residues 277-304 and 277-314 within the RBR linker were deleted (FoxP3^ΔN’^ and FoxP3^ΔN”^ in Figure S2A). Both constructs displayed preferential binding to IR-FKHM^4g^ as with FoxP3^ΔN^ (Figure S2B). Crystals of FoxP3^ΔN’^ and FoxP3^ΔN”^ diffracted to 4.0 Å and 3.1 Å, respectively (Table S1). As expected, our attempt to solve the structure using the swap-dimeric FoxP3^forkhead^ as the molecular replacement template failed. However, using non-swap monomeric structures of forkhead from other TFs (such as FoxN1 (Newman et al., 2020)), we identified unambiguous solutions and obtained final models (See Methods and Table S1).

The structures of FoxP3^ΔN’^ and FoxP3^ΔN”^ were similar and revealed several novel features of FoxP3. First, both FoxP3^ΔN’^ and FoxP3^ΔN”^ form the winged-helix, non-swap conformation, rather than the swap dimer (Figure 1E, Figure S2C). As with other forkhead TFs with the winged-helix fold (Dai et al., 2021), FoxP3 inserts the signature helix 3 (H3) into the major groove forming a sequence-specific interaction with DNA, while the wing 1 region forms additional contact with the DNA phosphate backbone. The potential source of the discrepancy between our structure versus the previous swap dimer structure and the functional implications of the new structure will be discussed in detail in Figure 2. Second, the structure also revealed that FoxP3^ΔN^ binds IR-FKHM^4g^ as a head-to-head (H-H) dimer, where each monomer occupies individual FKHM (Figure 1E). In this configuration, the forkhead dimer binds one side of the DNA, occupying two consecutive major grooves. DNA is bent by ∼25° (Figure 1E), although the degree of bending differs slightly between the FoxP3^ΔN’^ and FoxP3^ΔN”^ structures (Figure S2D). H-H dimerization of FoxP3 will be discussed in detail in Figures 3 and 4. Third, whereas structures of forkhead and part of RBR (residues 322-336) were resolved, the rest of RBR (residues 262-321), CC and ZF were not seen in either crystal structure. SDS-PAGE analysis, however, showed that the protein was intact in the crystal (Figure S2E), suggesting that RBR (262-321), ZF and CC are flexible or adopt heterogeneous conformations in the crystal. Note that ZF made a significant contribution to FoxP3–DNA interaction as evidenced by the negative impact of the ZF mutation K215A on DNA binding (Figure S2F). However, isolated ZF-CC did not bind DNA (Figure S2G). Based on these observations, we propose an overall architecture of FoxP3–DNA complex, where the H-H dimer of FoxP3^RBR-forkhead^ forms the primary contact with DNA, while two ZFs flexibly tethered through CC and RBR form secondary contacts with nearby sites on DNA without a fixed location relative to FoxP3^RBR-forkhead^ (Figure S2H).

**Figure 2.**
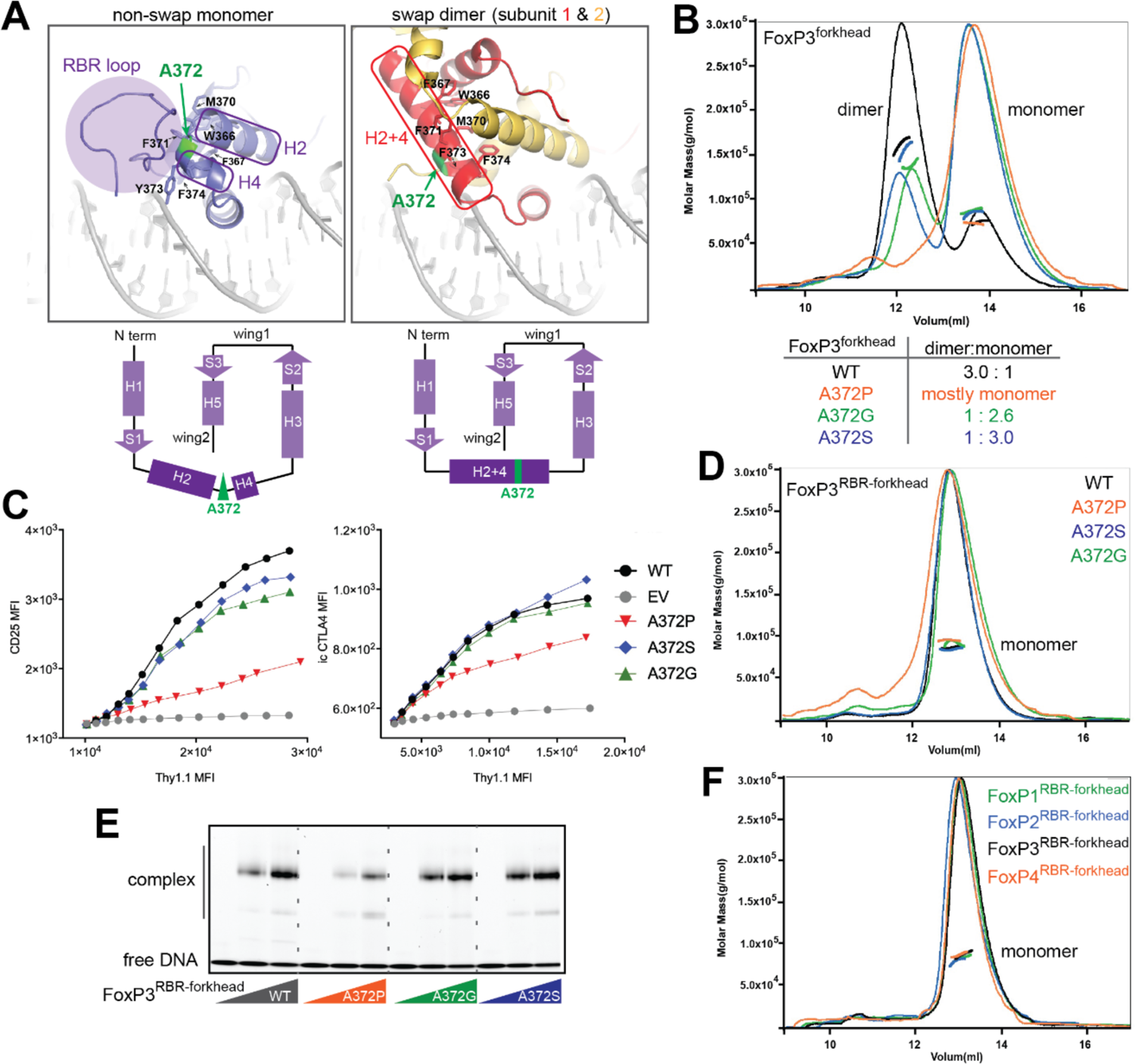
FoxP3 folds into the non-swap monomer in the native domain architecture. **See also Figure S3**. A. Structural comparison between non-swap monomer (our structure) of FoxP3 and swap dimer (previous structure, PDB: 3QRF). Hydrophobic residues (W366, F367, M370, F371, F373 and F374) lining the swap dimerization interface (right) are shown in sticks. Note that these residues in the non-swap monomer are folded to form a hydrophobic core protected by the RBR loop (left). Bottom: the two structures have identical secondary structure topology, except for helix 2 (H2) and helix 4 (H4), which are merged into one helix (H2+4) in the swap dimer. The residue A372 (green) is located in the junction between H2 and H4. B. SEC-MALS of NusA-tagged FoxP3^forkhead^ with and without mutations in A372. Below: dimer-to-monomer ratio was compared using peak intensities. C. FoxP3 cellular activity of swap-suppressive mutants, as measured by FACS. CD4^+^ T cells were retrovirally transduced to express FoxP3 with and without mutations in A372. Transcriptional activity of FoxP3 was analyzed by intracellular (i.c.) staining of CTLA4 or cell surface staining of CD25. FoxP3 expression was measured by Thy 1.1, which is under the control of IRES from the bicistronic mRNA expressing FoxP3. D. SEC-MALS of NusA-tagged FoxP3^RBR-forkhead^ with and without mutations in A372, all showing only one population corresponding to monomers. E. EMSA of NusA-tagged FoxP3^RBR-forkhead^ (0, 0.4 and 0.8 μM) with and without mutations in A372. DNA with IR-FKHM^4g^ was used. F. SEC-MALS of NusA-tagged FoxP1^RBR-forkhead^, FoxP2^RBR-forkhead^, FoxP3^RBR-forkhead^ and FoxP4^RBR-forkhead^, all showing only one population corresponding to monomers. Data in (B-F) are representative of at least three independent experiments.

**Figure 3.**
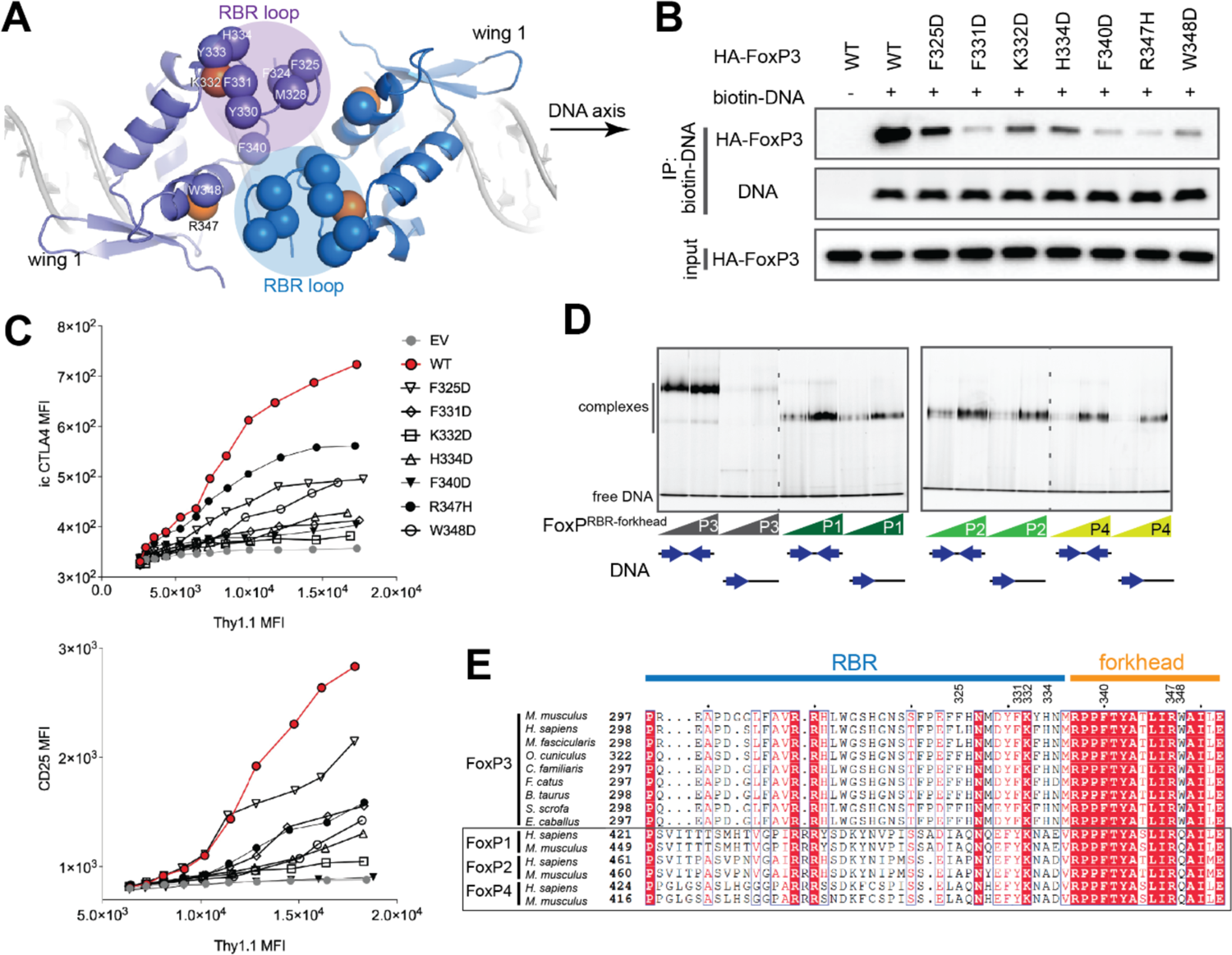
RBR loop-mediated H-H dimerization is important for and unique to FoxP3. **See also Figure S4**. A. Top view of the FoxP3 H-H dimer. The RBR loop forms the interface through the RBR– RBR and RBR–forkhead interactions. Hydrophobic residues (C*α*) at or near the interface are shown in blue or purple spheres. K332 and R347 are shown in orange spheres. B. DNA binding activity of H-H interface mutants. Biotinylated DNA with IR-FKHM^4g^ was used to pull-down FoxP3 (WT or mutants) ectopically expressed in 293T cells. C. FoxP3 cellular activity of H-H interface mutants. Experiments were performed as in Figure 2C. D. EMSA of NusA-tagged FoxP1-4 (0.4 and 0.8 μM) using DNA oligos (0.2 μM) with IR-FKHM^4g^ or single FKHM. All proteins were RBR-forkhead domains fused with NusA. E. Sequence alignment of FoxP3 orthologs and paralogs in the FoxP family. Data in (B-D) are representative of at least three independent experiments.

**Figure 4.**
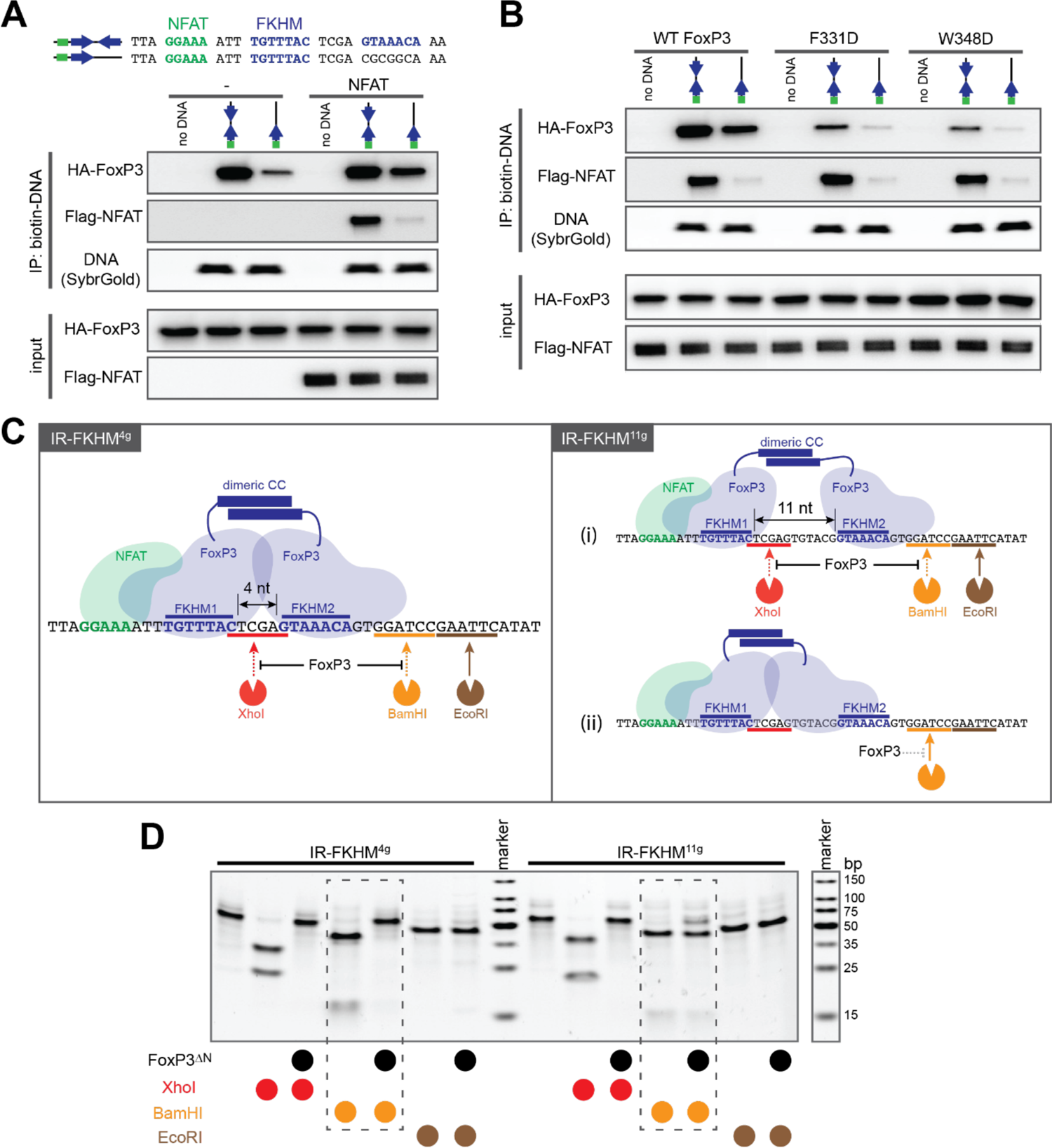
FoxP3 binds DNA as a H-H dimer, independent of DNA sequence. See also Figure S5. A. Biotin-DNA pull-down to monitor FoxP3*−*DNA interaction in the presence and absence of NFAT. DNA oligos with IR-FKHM^4g^ and single FKHM were compared. Full-length HA-tagged FoxP3 and FLAG-tagged NFAT were ectopically expressed in 293T cells and were subject to pull-down. B. Biotin-DNA pull-down to monitor FoxP3*−*DNA interaction in the presence and absence of NFAT. FoxP3 WT and two H-H interface mutants (F331D and W348D, see Figure 3) were compared. C. Schematic of the restriction enzyme protection assay. Left: H-H dimerization of FoxP3 on IR-FKHM^4g^ (47 bp) should protect the DNA from cleavage by XhoI and BamHI, but not by EcoRI. Right: two potential binding modes of FoxP3 for IR-FKHM^11g^ (54 bp), where the difference is in the accessibility of second FKHM (FKHM2), which can be examined by the BamHI site protection. D. Restriction enzyme protection assay with Foxp3^ΔN^ in the presence of NFAT. Note that higher FoxP3^ΔN^ concentration was used for IR-FKHM^11g^ (3.2 μM) than for IR-FKHM^4g^ (1.6 μM) so that all DNA is fully occupied by FoxP3 during footprint analysis. NFAT (0.4 μM) and DNA (0.2 μM) were used.

### FoxP3 forkhead exists in the non-swap monomeric conformation in the presence of RBR

Our finding that the forkhead domain within FoxP3^ΔN^ forms the non-swap monomeric conformation contradicts the previous report that isolated FoxP3^forkhead^ exists as a domain-swap dimer. We thus investigated the potential source of the discrepancy and which of the two is functionally relevant. The primary difference between the swap dimer and non-swap monomer is in the helix 2 (H2) and helix 4 (H4), which in the non-swap monomer are separated by a turn at residue A372 (Figure 2A, left, see both the topology diagram and structure). In contrast, in the swap dimer, H2 and H4 are merged into a single extended helix (H2+4) (Bandukwala et al., 2011) (Figure 2A, right, see both the topology diagram and structure). Intriguingly, the swap dimer interface on H2+4 contains a patch of hydrophobic residues (W366, F367, M370, F371, Y373 and F374), which in the non-swap conformation are buried within the folded forkhead core or protected by RBR (Figure 2A). This suggests that RBR may be responsible for stabilizing the non-swap conformation by protecting otherwise solvent exposed hydrophobic residues. In other words, deletion of RBR and subsequent exposure of hydrophobic residues may drive the swap dimerization. This view is supported by our observation that FoxP3 forkhead was monomeric in the presence of RBR (as in FoxP3^ΔN^ and FoxP3^RBR-forkhead^), while isolated FoxP3^forkhead^ without RBR formed a swap dimer (Figure 1B).

To further examine which of the two conformations is physiologically relevant, we employed a protein engineering approach with the focus on residue 372 at the junction of H2 and H4 (Figure 2A). This is because this position is unique in its potential to switch the forkhead conformation between swap and non-swap structures, as previously noted (Bandukwala et al., 2011). We hypothesized that replacement of Ala at 372 by an amino acid with low helix propensity, such as Pro, Gly or Ser, would force separation of H2 and H4 and convert FoxP3^forkhead^ to the monomeric conformation. As predicted, A372P, A372G and A372S all biased isolated FoxP3^forkhead^ from dimeric to largely monomeric state, albeit to varying degrees (Figure 2B). Protein crosslinking analysis also suggested that all three mutations suppressed swap dimerization (Figure S3A). We next examined the impact of the A372 mutations on the transcriptional activity of FoxP3, as measured by the levels of CTLA4 and CD25 (two Treg markers) upon retroviral expression of FoxP3 in CD4^+^ T cells. If FoxP3 functions require the swap-dimeric structure, all three mutations of A372 should result in loss of function. Arguing against this idea, we found that A372G and A372S had little to mild impact on the transcriptional activity of FoxP3 (Figure 2C). A372P, on the other hand, partially impaired FoxP3 function (Figure 2C), as previously reported (Bandukwala et al., 2011). This negative impact of A372P, however, appeared to be independent of its ability to suppress swap dimerization; introduction of A372P in FoxP3^RBR-forkhead^, which already existed as a non-swap monomer, did not alter its monomeric state (Figure 2D), but negatively affected its DNA affinity (Figure 2E) and drastically altered the protein melting curve (Figure S3B). Such negative impact was not observed with A372G and A372S (Figures 2E and S3B), further suggesting that the functional impairment by A372P is due to its effect on DNA affinity and protein thermal stability rather than its swap-suppressive property. Collectively, the results for A372G and A372S argue against the model that swap dimerization is required for FoxP3 transcriptional activity.

We next asked whether the impact of RBR on forkhead folding is conserved in other FoxP TFs, which also contain Ala at position equivalent to 372 (Figure S3C) and were also reported to form domain-swap dimers when expressing FKHs in isolation (Chu et al., 2011; Stroud et al., 2006). FoxP1, FoxP2 and FoxP4 all contain RBR-like linkers between CC and FKH. We expressed FoxP1/2/4 proteins equivalent to the FoxP3^RBR-forkhead^ construct and found that FoxP1/2/4 proteins were monomeric when purified with RBR (Figure 2F). This observation suggests that the role of RBR or RBR-like linkers in stabilizing the non-swap conformation is conserved in all FoxP TFs. They further support the notion that the non-swap conformation is the physiologically relevant form for all four members of FoxP TFs.

### Head-to-head (H-H) dimerization of forkhead is unique to FoxP3 and is important for DNA binding

Our structure shows that individual FoxP3 forkhead folds into a monomer, but this monomer forms a H-H dimer when bound to DNA containing IR-FKHM^4g^ (Figure 1E). H-H dimerization is mediated by part of RBR (residues 321-336, to be referred to as the RBR loop), which interacts with the RBR loop and forkhead of the other subunit (Figure 3A). We also note that electron density for the RBR loop was not as well defined as other parts of the protein (Figure S4A), suggesting conformational flexibility in the interface. Intriguingly, residues located within the RBR loop and near the interface are highly hydrophobic (Figure 3A), suggesting that H-H dimerization is driven by a collection of hydrophobic interactions, reminiscent of “fuzzy” interactions shown for other TFs (Pricer et al., 2017; Tuttle et al., 2021). It is noteworthy that the RBR loop is also involved in crystallographic packing (Figures S4A, S4B), likely involving similarly fuzzy, hydrophobic interactions. Thus, the hydrophobic nature of the RBR loop may enable multiple types of protein-protein interactions beyond H-H dimerization (to be discussed in Figure 6).

To examine whether the H-H dimerization is simply a result of binding IR-FKHM^4g^ or accounts for FoxP3’s preference for IR-FKHM^4g^, we introduced single-point mutations in or near the H-H dimeric interface in both RBR and forkhead. These include aforementioned hydrophobic residues and basic residues (K332 and R347) that are in the position to favorably interact with aromatic residues (Figure 3A). Note that R347H is an IPEX mutation (Gambineri et al., 2008).

Consistent with the view that H-H dimerization is the cause, rather than the consequence, of IR-FKHM^4g^ binding, the mutations impaired FoxP3’s affinity for DNA containing IR-FKHM^4g^, as measured by biotinylated DNA pull-down (Figure 3B). Some residues in the RBR loop, such as F331 and H334, are not located at the interface but still played important roles, suggesting that these residues likely play a role in shaping the RBR loop and thus are indirectly involved in H-H dimerization. The interface mutations also impaired the transcriptional activity of FoxP3 (Figure 3C). These results suggest that H-H dimerization is important for both DNA binding and FoxP3’s transcriptional activity.

Given the importance of H-H dimerization for FoxP3, we next asked whether H-H dimerization also occurs with other FoxP TFs. This was plausible given the importance of RBR in FoxP3 H-H dimerization and the common role of RBR/RBR-like linker in stabilizing the non-swap conformation in all four FoxP TFs (Figure 2F). To examine the possibility of H-H dimerization, we compared FoxP^RBR-forkhead^ binding to IR-FKHM versus sFKHM, where H-H dimerization would manifest in preferential binding of IR-FKHM. We used FoxP^RBR-forkhead^, instead of the FoxP^ΔN^ construct due to the lack of solubility of FoxP1^ΔN^, FoxP2^ΔN^ and FoxP4^ΔN^. Note that both FoxP3^ΔN^ and FoxP3^RBR-forkhead^ preferentially bound IR-FKHM^4g^ (Figure 3D, S1D and S4C). In contrast, FoxP1/2/4 equally bound IR-FKHM^4g^ and sFKHM (Figure 3D), and this is independent of the IR-FKHM gap size (Figure S4C). Additionally, the native gel migration rate of the FoxP1/2/4 complexes were similar, either on IR-FKHM^4g^ or sFKHM – another indication that they bound both DNAs as a monomer (Figure 3D). These results suggest that H-H dimerization does not occur to other FoxP TFs. Perhaps reflecting this difference in the ability to form a H-H dimer, the RBR sequence of FoxP3 differs from the equivalent linker in other FoxP TFs (Figure 3E). Altogether, our results suggest that H-H dimerization is an important and a unique feature of FoxP3 and imparts distinct DNA specificity to FoxP3.

### H-H dimerization is important for DNA binding, regardless of DNA sequence

We next asked whether H-H dimerization of FoxP3 is limited to IR-FKHM^4g^ or whether this also occurs when FoxP3 binds suboptimal, low-affinity sequences. This is an important question because many TFs are known to bind suboptimal sequences in cells (Pfeifer et al., 1987; Segal et al., 2008), likely driven by cooperative binding with cofactors (Reiter et al., 2017). In fact, FoxP3 is also known to function together with other cofactors (Kwon et al., 2017; Rudra et al., 2012; Wu et al., 2006), many of which have their own DBDs and were hypothesized to drive FoxP3–DNA interaction in cells (Samstein et al., 2012; Wu et al., 2006). One of the best-characterized cofactors for FoxP3 is NFAT, whose DBD (Rel Homology Region, RHR) interacts directly with FoxP3 forkhead as shown by the crystal structure of the complex (Bandukwala et al., 2011). Note that this structure was determined with the swap dimeric forkhead, but the NFAT^RHR^ interface is nearly identical in both the swap and non-swap conformations, making both conformations compatible with NFAT binding (Figure S5A). As such, FoxP3^ΔN^, which is in the non-swap conformation, also showed cooperative DNA binding with NFAT when the NFAT-binding site is 3 nt upstream of FKHM, the requirement for NFAT–FoxP3 interaction (Figure 4A). This cooperative relationship was more evident with sFKHM than with IR-FKHM^4g^, as evidenced by the positive effect of NFAT on FoxP3 binding to sFKHM and the lack of such effect with IR-FKHM^4g^ (Figure 4A). This suggests that NFAT can drive FoxP3 binding to suboptimal DNA sequences, but its role could be minimal for optimal DNA, such as IR-FKHM^4g^. When H-H interface mutations (F331D and W348D, chosen from Figure 3) were introduced to FoxP3, DNA affinity was reduced for both IR-FKHM^4g^ and single FKHM, with or without NFAT (Figure 4B). This suggests that H-H dimerization is important even when FoxP3 binds suboptimal DNA driven by the cofactor NFAT.

To more directly examine whether FoxP3 forms the H-H dimer when it binds suboptimal DNA sequence, we examined FoxP3 footprint using a restriction enzyme protection assay. We chose IR-FKHM^11g^ as a suboptimal DNA because it allows for predicting two distinct binding modes. In one mode, two FoxP3 molecules bridged by CC may individually occupy two FKHMs separated by the 11 nt gap without forming a H-H contact (Figure 4C, right panel (i)). In the alternative mode, FoxP3 may bind IR-FKHM^11g^ as a H-H dimer, occupying only one of the FKHMs – most likely FKHM near the NFAT site (FKHM1) – and the suboptimal sequence 4 nt away from FKHM1, leaving the second FKHM (FKHM2) unoccupied (Figure 4C, right panel (ii)). To distinguish between the two modes, we placed the BamHI restriction site near FKHM2 to examine the occupancy of FKHM2 (Figure 4C). The XhoI and EcoRI sites were also introduced to report FKHM1 occupancy and DNA binding specificity, respectively.

We first tested an equivalent construct with IR-FKHM^4g^. Under the condition where FoxP3^ΔN^ occupies DNA in a sequence-specific manner (*i.e.* the XhoI site was fully protected while the EcoRI site was fully accessible), we found that the BamHI site was protected by FoxP3 (Figure 4D, left half), consistent with the crystal structure where both FKHM1 and FKHM2 are occupied by FoxP3 H-H dimer. In contrast, the BamHI site of IR-FKHM^11g^ was largely accessible even though DNA was fully occupied by FoxP3 (as evidenced by the XhoI site protection) (Figure 4D, right half). Under the equivalent condition, FoxP3^forkhead^ (A372S), which does not form a H-H dimer, showed no difference between IR-FKHM^4g^ and IR-FKHM^11g^ in the BamHI site protection (Figure S5B). These results suggest that FoxP3^ΔN^ maintains the H-H dimeric structure when binding IR-FKHM^11g^ as well as IR-FKHM^4g^. Together with the data in Figure 4B, they further support the notion that FoxP3 binds DNA as a H-H dimer, regardless of the DNA sequence.

### H-H dimerization enables FoxP3 to recognize diverse DNA sequences

Our data above suggested that H-H dimerization is likely to be an important mode of DNA binding for FoxP3 and this greatly influences its DNA sequence specificity, as evidenced by the strong preference for IR-FKHM^4g^ in vitro. To assess the impact of H-H dimerization on FoxP3– DNA interaction in Treg cells, we investigated previously reported FoxP3 ChIP-seq data for potential enrichment of the IR-FKHM^4g^ sequence (Kitagawa et al., 2017; Samstein et al., 2012). *De novo* motif analysis of 5000 consensus FoxP3 ChIP-seq peaks did not identify IR-FKHM^4g^ (Figure 5A), although single FKHM was identified in these peaks (P-value<10^-9^) (Figure 5A). As an alternative strategy, we separately analyzed 548 FoxP3-bound peaks that contain an FKHM and counted the number of instances of an inverted FKHM repeats with a gap size ranging from 1 to 21 nt; no enrichment of 4 nt spacing was observed (Figure 5B). One explanation for this lack of IR-FKHM^4g^ enrichment could be that FoxP3, like other TFs (Nakagawa et al., 2013), utilizes many distinct suboptimal sequences in the presence of cofactors. H-H dimerization of FoxP3 may further increase the diversity of such suboptimal sequences it can bind, as shown with other dimeric TFs (Arnett et al., 2010; Jiang et al., 2019).

**Figure 5.**
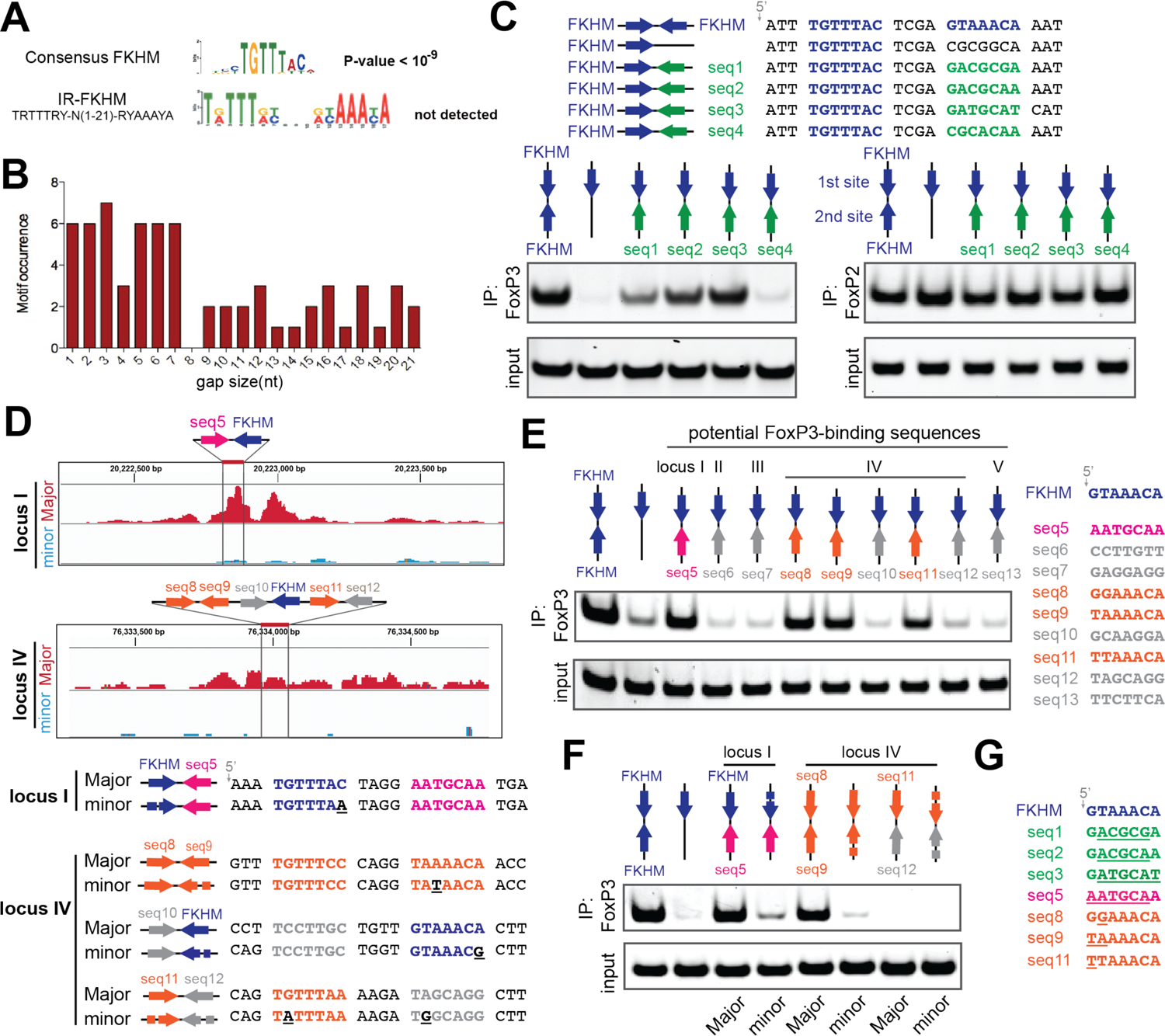
H-H dimerization enables FoxP3 to recognize diverse sequences. See also Figure S6. A. *De novo* motif analysis of FoxP3 ChIP-seq (n=5000) sequences. The canonical FKHD motif was enriched (P-value<10^-9^), but IR-FKHM motif was not detected for any of the gap sizes tested (1-21 nt). A more relaxed FKHM sequence (TRTTRY; R and Y indicate purine and pyrimidine, respectively) was used to be inclusive. B. Occurrence of IR-FKHM, iteratively testing gap sizes of 1 to 21 nt, of FoxP3 ChIP-seq sequences that contain an FKHM (n=548). Motif occurrences were counted if p-value<10^-5^ and score greater than 12. C. FoxP3 interaction with DNA harboring IR-FKHM^4g^ and FKHM paired with non-canonical motifs (seq1-4). Purified MBP-FoxP3^ΔN^ was mixed with DNA oligos and was subjected to MBP pull-down, followed by native PAGE analysis of co-purified DNA. DNA was visualized by Sybr Gold stain. Right: MBP-FoxP2^ΔN^ was used for comparison. D. FoxP3 Cut&Run intensity showing allelic imbalance of FoxP3 occupancy in loci I and IV. Major and minor alleles indicate alleles with greater and lesser FoxP3 occupancy, respectively. Below: Major and minor allele sequences with their differences highlighted with underscores in the minor sequences. E. FoxP3 interaction with DNA harboring potential FoxP3-binding sequences paired with FKHM. These sequences were chosen from five loci in the Major allele sequence where mutations were associated with reduced FoxP3 occupancy. F. FoxP3 interaction with DNA harboring natural sequences from loci I and IV (see D). G. Alignment of FoxP3-compatible sequences examined in this figure. Sequence deviation from FKHM was highlighted with underscore. Data in (C, E and F) are representative of at least three independent experiments.

To examine the possibility of diverse sequence recognition by the FoxP3 H-H dimer, we first examined FoxP3 binding to four non-consensus motifs (seq1-4) that were previously shown to bind other forkhead TFs (Nakagawa et al., 2013; Rogers et al., 2019). These non-consensus sequences were paired with FKHM in the inverted orientation with a 4 nt gap (Figure 5C, top). FoxP3 bound FKHM paired with seq1/2/3 significantly more efficiently than sFKHM (Figure 5C, bottom left). In comparison, FoxP2 showed similar binding to all DNA regardless of the sequence paired with FKHM (Figure 5C, bottom right). When seq1-3 were paired with themselves instead of FKHM, FoxP3 binding was undetectable (Figure S6C), indicating that at least one FKHM is necessary. Similar results were obtained using FoxP3 by EMSA (Figures S6A and S6B). These results suggest that strong anchoring of one FoxP3 monomer at FKHM relaxes the sequence requirement for the second site, allowing FoxP3 to bind other sequences besides IR-FKHM^4g^.

We next examined physiological target sites of FoxP3. A previous Cut&Run analysis (van der Veeken et al., 2020) identified sequence-specific FoxP3-binding sites based on the observation that allelic variations in the FKHM sequence between two evolutionary distant mouse genomes were accompanied by reduced FoxP3 occupancy (see Figure 5D for examples; “Major” and “minor” alleles indicate those with greater and lesser FoxP3 occupancy, respectively). We asked whether these sites contain FoxP3-compatible sequences as the second binding site. Sequences paired with FKHM in five loci with allelic imbalance in FoxP3 occupancy were tested. Locus IV had several FKHM-like sequences (seq8, seq9 and seq11; Figure 5D), which were also included in our analysis. Among the 9 sequences tested, four sequences (seq5/8/9/11) efficiently bound FoxP3 when paired with FKHM (Figure 5E), but not when paired with themselves (Figure S6D). Thus, loci I and IV have suboptimal, but nevertheless FoxP3-compatible sequences (seq5/8/9/11) paired with each other or with FKHM. However, loci II, III and V do not contain FoxP3-compatible sequences paired with FKHM, suggesting that FoxP3 may bind elsewhere within these loci without FKHM.

We further examined loci I and IV. As expected, FoxP3 binding to FKHM-seq5 of locus I was markedly reduced upon introduction of the mutation in FKHM (as in the minor allele) (Figure 5F), consistent with the view that FKHM-seq5 is responsible for FoxP3 binding at locus I. In contrast to locus I, locus IV contains multiple potential FoxP3 binding sites: seq8 paired with seq9, seq10 with FKHM, and seq11 with seq12 (Figure 5D). Of these, seq8-seq9 is the only site where FoxP3-compatible sequences are paired. Accordingly, only seq8-seq9 showed strong FoxP3 binding, while seq10-FKHM and seq11-seq12 did not (Figure 5E and 5F). Furthermore, sequence variation in seq9 (as in the minor allele) markedly reduced FoxP3 binding (Figure 5F), supporting the view that seq8-seq9 significantly contributes to locus IV binding. Note that both seq8 and seq9 are FKHM-like sequences, but neither is strong enough to recruit FoxP3 alone or when paired with itself (Figure S6D). This suggests that certain combinations of motifs can have a non-additive effect and pairing of suboptimal sequences with the optimal FKHM is not obligatory.

We note that the non-consensus sequences that we identified here do not display any obvious patterns (Figure 5G), which may explain why the second FoxP3 site could not be detected from the global motif analysis. These non-consensus sequences are also likely to be a small subset of a possibly much larger pool of sequences FoxP3 binds in the presence of cofactors in cells, as evidenced by the lack of enrichment of these compatible sequences, relative to non-compatible sequences, within the ChIP-seq peaks (Figure S6E). Nevertheless, our findings suggest that H-H dimerization significantly alters FoxP3 DNA specificity and enables FoxP3 to recognize diverse sequences beyond IR-FKHM^4g^.

### H-H dimerization is important for Runx1 binding

Given that H-H dimerization is largely mediated by RBR and that RBR was previously shown to play important roles in recruiting Runx1 (Ono et al., 2007), we next asked what effect H-H dimerization has on Runx1 binding. Our structure showed that H-H dimerization creates a large hydrophobic surface opposite from the DNA binding surface, which mediates crystallographic packing (Figure S4B). Given that FoxP3-binding region of Runx1 (residue 371-451) is highly hydrophobic (Figure 6A), we asked whether this hydrophobic surface of FoxP3 could serve as a cofactor docking site for Runx1.

Using purified recombinant Runx1 (Runx1^ΔN^, residue 371-451) and FoxP3^RBR-forkhead^, we confirmed that their interaction is direct (Figure 6B). We also found that this interaction was enhanced in the presence of DNA with IR-FKHM^4g^ (Figure 6B), the condition that promotes H-H dimerization. The Runx1–FoxP3 interaction was not enhanced by DNA harboring sFKHM, further supporting the notion that H-H dimerization is important for Runx1 binding. Since Runx1^ΔN^ does not contain a DBD, the observed IR-FKHM^4g^-dependent interaction between Runx1 and FoxP3 cannot be due to DNA-mediated bridging. Using full-length Runx1 and full-length FoxP3 overexpressed in 293T cells, we also confirmed that their association is dependent on nucleic acids, as the benzonase treatment significantly impaired their co-immunoprecipitation (co-IP) (Figure 6C). Consistent with the idea that Runx1 binding is facilitated by DNA-dependent H-H dimerization, H-H interface mutations also reduced their co-IP (Figure 6D). Together, these data suggest that FoxP3 uniquely harnesses its H-H dimerization capability not only to alter DNA sequence specificity, but also to recruit Runx1 and to coordinate it with DNA binding.

**Figure 6.**
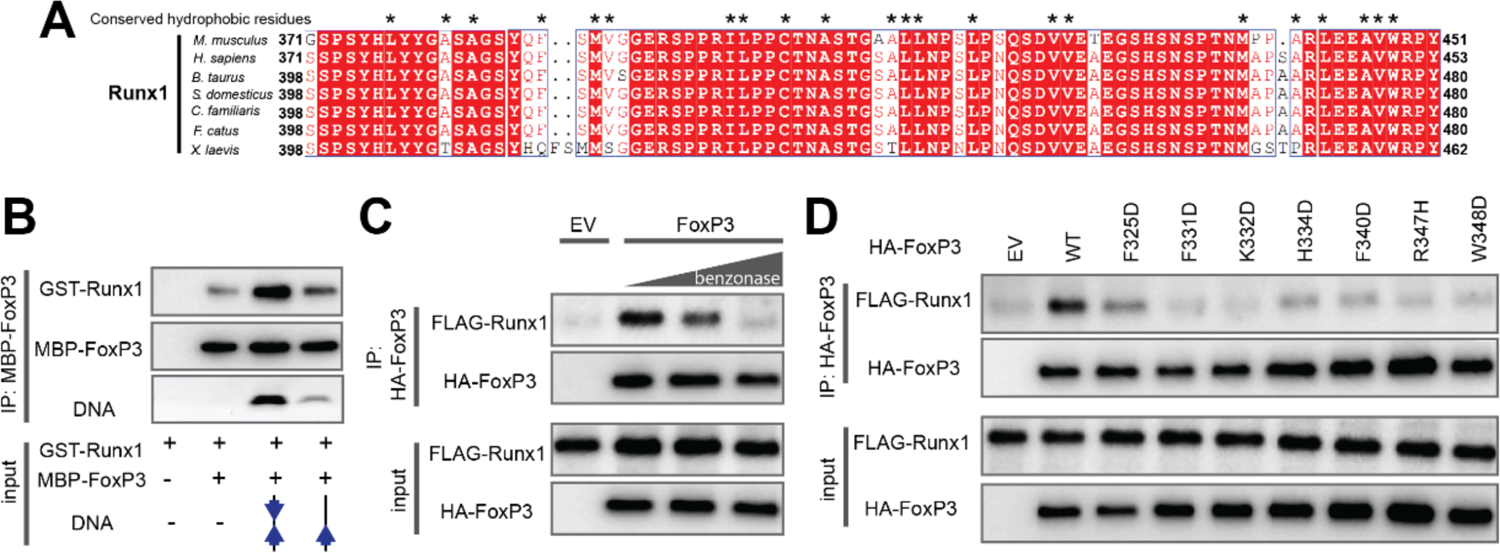
H-H dimerization is also important for Runx1 binding. A. Sequence alignment of Runx1 orthologs showing conserved hydrophobic residues (*) in its C-terminal tail that binds FoxP3. B. FoxP3 interaction with Runx1 using purified proteins. MBP-tagged FoxP3^RBR-forkhead^ and GST-tagged Runx1 (residue 371-451) was purified from *E. coli* and was subjected to MBP pull-down in the presence and absence of DNA harboring IR-FKHM^4g^ or single FKHM. C. FoxP3 interaction with Runx1 from 293T cells. HA-tagged full-length FoxP3 and FLAG-tagged full-length Runx1 were separately expressed in 293T cells. Cell lysates were mixed and were subjected to anti-HA immunoprecipitation (IP). The increasing concentrations of benzonase was used to examine the effect of cellular DNA on FoxP3– Runx1 interaction. D. FoxP3 interaction with Runx1 from 293T cells. H-H interface mutations were compared to WT FoxP3. Experiments were performed as in (C), except no benzonase was used in all conditions. Data in (B-D) are representative of at least three independent experiments.

### IPEX mutation R337Q induces swap dimerization and impairs DNA and Runx1 binding

Given our findings supporting the importance of H-H dimerization and non-swap conformation, we asked whether some IPEX mutations are caused by errors in these features. Note that the non-swap conformation is a precondition for H-H dimerization, and thus, any mutation that alters non-swap folding would impair all aspects of FoxP3 function including H-H dimerization, DNA binding and Runx1 binding. Among the IPEX-associated mutations of *FOXP3*, R337Q attracted our attention as it is located near the end of RBR (Rubio-Cabezas et al., 2009) (Figure 1A), which is involved in both stabilization of the non-swap conformation and H-H dimerization. Additionally, R337 directly contacts DNA in our non-swap structure, but is far away from DNA in the swap-dimeric structure (Figure 7A), a feature that could further help distinguish between the two forkhead conformations.

**Figure 7.**
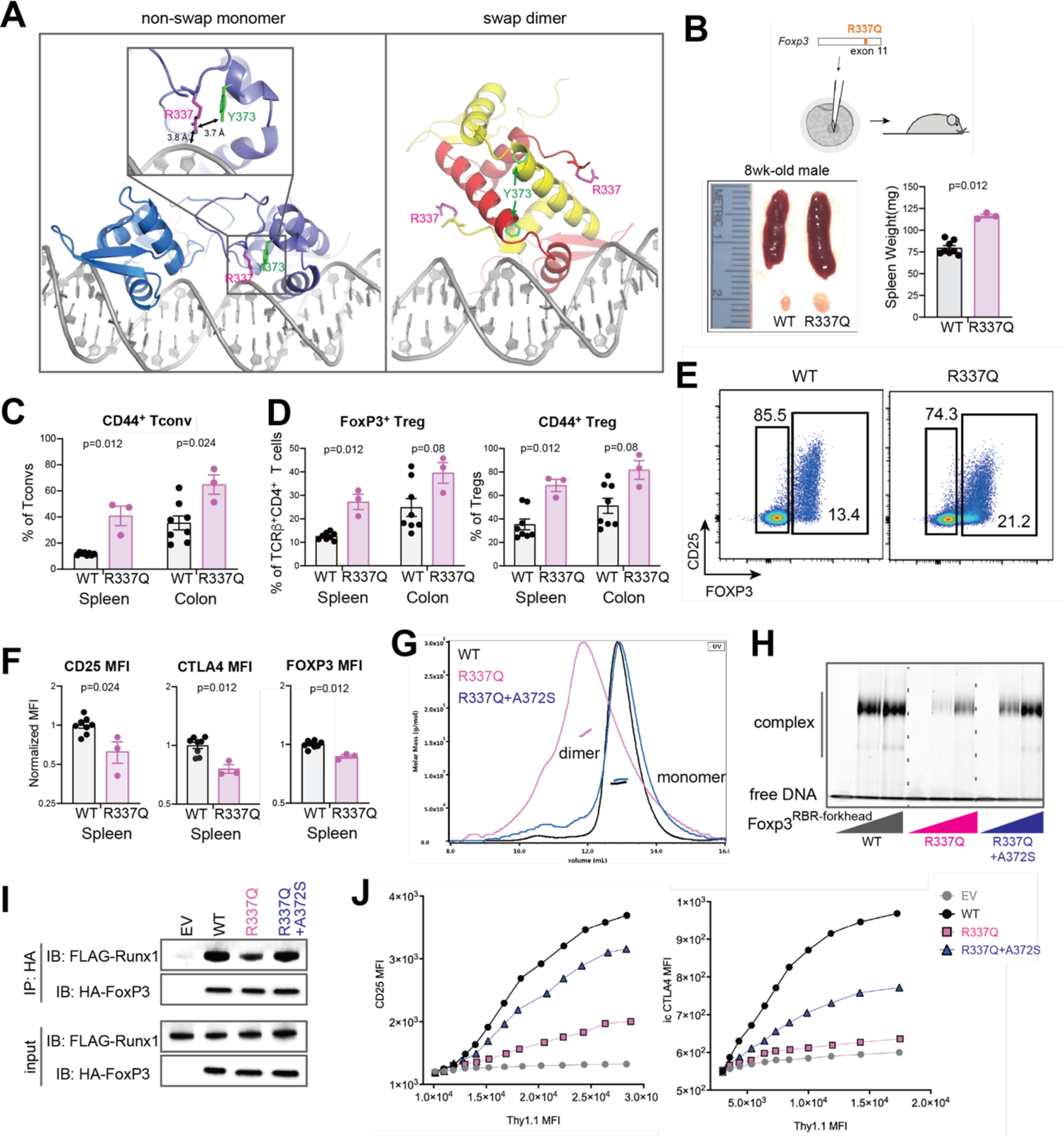
IPEX mutation R337Q induces swap dimerization, impairing DNA binding and Runx1 recruitment. **See also Figure S7**. A. Location of R337 in the crystal structures of swap dimeric and non-swap monomeric FoxP3. R337 interacts with DNA and Y373 only in the non-swap conformation. B. Abnormal immune homeostasis in 8-week-old *R337Q* male mice, generated by CRISPR mutagenesis of *Foxp3*. Bottom left: a representative picture comparing the spleens and inguinal lymph nodes of *R337Q* mutant mice and their WT littermates. Bottom right: spleen weights. C. Fraction of activated (CD44^+^) conventional T (Tconv) cells among CD4^+^ T cells in the spleen and the colonic lamina propria of WT and R337Q mutant mice. D. Foxp3^+^ Treg in the spleen and the colonic lamina propria of *R337Q* mutant mice in comparison to their WT littermates. Left: their fraction among TCR*β*^+^CD4^+^ T cells. Right: their activation status assessed by the marker CD44^+^. E. CD25/FoxP3 cytometry plots of spleen TCR*β*^+^CD4^+^ T cells from WT and *R337Q* mutant mice. F. Quantification of CD25, CTLA4 and FoxP3 by flow cytometry (MFI) in TCR*β*^+^CD4^+^Foxp3^+^ spleen Tregs of *R337Q* mutant mice and WT littermates. G. SEC-MALS of NusA-tagged FoxP3^RBR-forkhead^ (WT, R337Q and R337Q+A372S) H. DNA binding activity of recombinant, purified FoxP3^RBR-forkhead^ (WT, R337Q and R337Q+A372S) as measured by EMSA. FoxP3 was tagged with NusA. DNA harbors IR-FKHM^4g^. I. Runx1 binding for full-length FoxP3 (WT, R337Q and R337Q+A372S) from 293T cells. J. Transcriptional activity of FoxP3 as measured by CD25 and CTLA4 levels. The effects of R337Q, either alone or in combination with the swap-suppressive mutation A372S, were examined. Data in (B-D, F) are presented as mean ± SEM. *P* values were obtained by Mann-Whitney test comparing WT and *R337Q*.. Data in (G-J) are representative of at least three independent experiments.

To better analyze the effect of R337Q in Treg cells *in vivo*, we generated by CRISPR/Cas9 germline mutagenesis a mutant mouse line carrying the R337Q mutation in the endogenous *Foxp3* locus (Figure 7B). Young male mice hemizygous for the mutation were viable, with no sign of wasting disease, or overt dermatitis or intestinal pathology, differing from full *FoxP3* deficiencies (Fontenot et al., 2003; Godfrey et al., 1991), although reduced weight gain and dermatitis did appear in older R337Q animals. In addition, they presented significant splenomegaly and lymphadenopathy (Figure 7B), and an increase in the level of activated (CD44^+^) T conventional (Tconv) cells (Figure 7C), suggesting a partial loss of Treg cell functionality. Importantly, the proportion of FoxP3^+^ Treg cells and their activated (CD44^+^) fraction also increased in R337Q mice (Figure 7D), a compensatory phenotype previously observed in partial FoxP3 deficiencies (Kwon et al., 2018; Van Gool et al., 2019). R337Q Treg cells also showed decreased levels of CD25 and FoxP3, as well as CTLA4, characteristic of FoxP3 hypomorphs (Figures 7E and 7F), all indicative of partial FoxP3 dysfunction. Thus, the R337Q mutation impairs Treg fitness, with a repercussion in the control of CD4^+^ T cell homeostasis, although not as radical as the typical *scurfy* phenotype usually seen with the complete *Foxp3* deficiencies.

We next introduced R337Q in recombinant FoxP3^RBR-forkhead^ and analyzed its impact on the biochemical properties of FoxP3. Intriguingly, FoxP3^RBR-forkhead^ with R337Q was purified as a dimer, which is in contrast to monomeric WT FoxP3^RBR-forkhead^ (Figure 7G). The R337Q dimer appeared to be a swap dimer, since addition of the swap-suppressive mutation A372S reverted R337Q FoxP3^RBR-forkhead^ to a monomer (Figure 7G). In line with the dependence of DNA binding on H-H dimerization, and thus the non-swap conformation, R337Q significantly reduced DNA affinity, while A372S partially restored it (Figure 7H and S7A). Similarly, Runx1 binding was also impaired by R337Q, but was largely restored by A372S (Figure 7I). Most importantly, the loss of FoxP3 transcriptional activity by R337Q was partially rescued by A372S (Figure 7J). The incomplete rescue likely reflects the role of R337 not only in stabilizing the non-swap conformation, but also in direct DNA binding, as our FoxP3 structure predicts (Figure 7A). Collectively, these data suggest that functional impairment by the R337Q mutation is largely mediated by its induction of swap dimerization, although its role in direct DNA binding also contributes.

The impact of R337Q on FoxP3 folding was unexpected as R337 is on the protein surface and folding errors are often associated with mutations in protein cores. Interestingly, R337 is in close contact with Y373, another surface residue mutated in IPEX patients (Y373V) (Tanaka et al., 2005) (Figure 7A). Recombinant Y373V FoxP3^RBR-forkhead^ protein displayed a broad elution profile in size-exclusion chromatogram with the peak molecular weight corresponding to tetramer (Figure S7B). This observation argues that Y373V also induces folding defects, albeit somewhat differently than R337Q. Consistent with aberrant formation of multimers, Y373V impaired FoxP3 transcriptional activity in T cells (Figure S7C). Together, these results suggest that FoxP3 may have unusual folding landscape that makes it prone to misfolding as a domain-swap dimer, and that swap-dimerization can naturally occur through loss-of-function IPEX mutations, such as R337Q.

## DISCUSSION

Towards our goal of understanding how FoxP3 utilizes the common DBD forkhead to function as a unique TF for Treg determinism, we characterized FoxP3 using a combination of structural biology, biochemistry and functional assays. Our study revealed that, unlike previous reports with isolated forkhead DBD (Bandukwala et al., 2011), FoxP3 forkhead folds into a non-swap monomer, and instead utilizes a novel type of H-H dimerization to acquire unique functions. H-H dimerization is required for DNA binding and this is mediated by the unique RBR linker region. This requirement for H-H dimerization distinguishes FoxP3 from other forkhead TFs characterized to date, including FoxP1/2/4, which can bind DNA as individual DBDs (Dai et al., 2021). As a result, FoxP3 reads DNA sequence spanning ∼18 bp, rather than 7 bp as monomeric forkhead does. While FoxP3 strongly favors inverted repeat of FKHM (IR-FKHM) over a single FKHM, it also allows diverse and distinct non-consensus sequences beyond IR-FKHM. This scope of diversity in FoxP3-compatible sequences is likely to expand further in the presence of cofactors in cells. FoxP3 H-H dimerization also influences its interaction with cofactors. We found that Runx1 binding is greatly facilitated by FoxP3 H-H dimerization, likely through the large hydrophobic patch formed upon dimerization. Since H-H dimerization is promoted by DNA binding, this Runx1 binding mechanism should enable FoxP3 to tightly coordinate DNA binding and Runx1 recruitment. Thus, a highly conserved DBD can diversify its function by gaining a multimerization capability, which may have implications beyond forkhead DBD.

A surprising finding was that RBR is not only important for H-H dimerization but also modifies forkhead DBD folding, providing an unusual example where an appendage domain alters folding of a highly conserved DBD. Given that it is a common practice to study isolated DBD for TF– DNA interactions, FoxP3 may serve as a cautionary tale for such approaches. In the absence of RBR, FoxP3 forkhead folds into the swap dimer as previously described (Bandukwala et al., 2011), while in the presence of RBR, it forms the canonical forkhead structure. Although the RBR-like linkers in FoxP1, 2, and 4 do not support H-H dimerization as for FoxP3, their roles in stabilizing the canonical forkhead structure was conserved across the FoxP family. Our data suggest that the canonical forkhead structure is likely the functional form as the swap dimer cannot engage with each other in a manner analogous to H-H dimerization and does not efficiently bind DNA and Runx1. We further showed that the IPEX mutation R337Q favors swap dimerization, compromising DNA and Runx1 binding and impairing the transcriptional activity, while a swap-suppressive mutation largely restored these functional impairments. These results altogether support that canonical non-swap conformation is the physiological form of FoxP3.

The unique dependence of FoxP3 folding on RBR and its sensitivity to surface residue mutations, such as R337Q, raise the question whether FoxP3 folding landscape is complex and folding errors are more common than just with R337Q. In other words, swap dimerization may not just be an *in vitro* artifact of expressing forkhead in isolation, but may readily occur in the presence of certain mutations. Supporting this view, we found that a mutation of another surface residue, Y373V, also leads to dramatic changes in protein multimerization – indicative of errors in protein folding – and loss of transcriptional activity in keeping with its association with IPEX. Based on these observations, we speculate that FoxP3 folding could be a novel therapeutic target; small molecules that either promote or destabilize the canonical monomeric forkhead structure or its H-H dimerization could be used to modulate Tregs in immunotherapy of cancer or autoimmunity.

## Acknowledgements

We thank members of the Hur lab for discussion and critical reading of the manuscript. This study was supported by NIH grants (R01AI154653 and R01AI111784 to S.H; AI150686 to C.B.; P30 CA008748 and R01 AI034206 to A.R.). J.L. was supported by an INSERM Poste d’Accueil and an Arthur Sachs scholarship, R.R. by NIH supplement AI116834-03S1. S.H. and A.R. are investigators at the Howard Hughes Medical Institute. X-ray diffraction data were collected at Advance Photon Source, beamline NECAT-24-ID-E.

---

**Figure S1.**
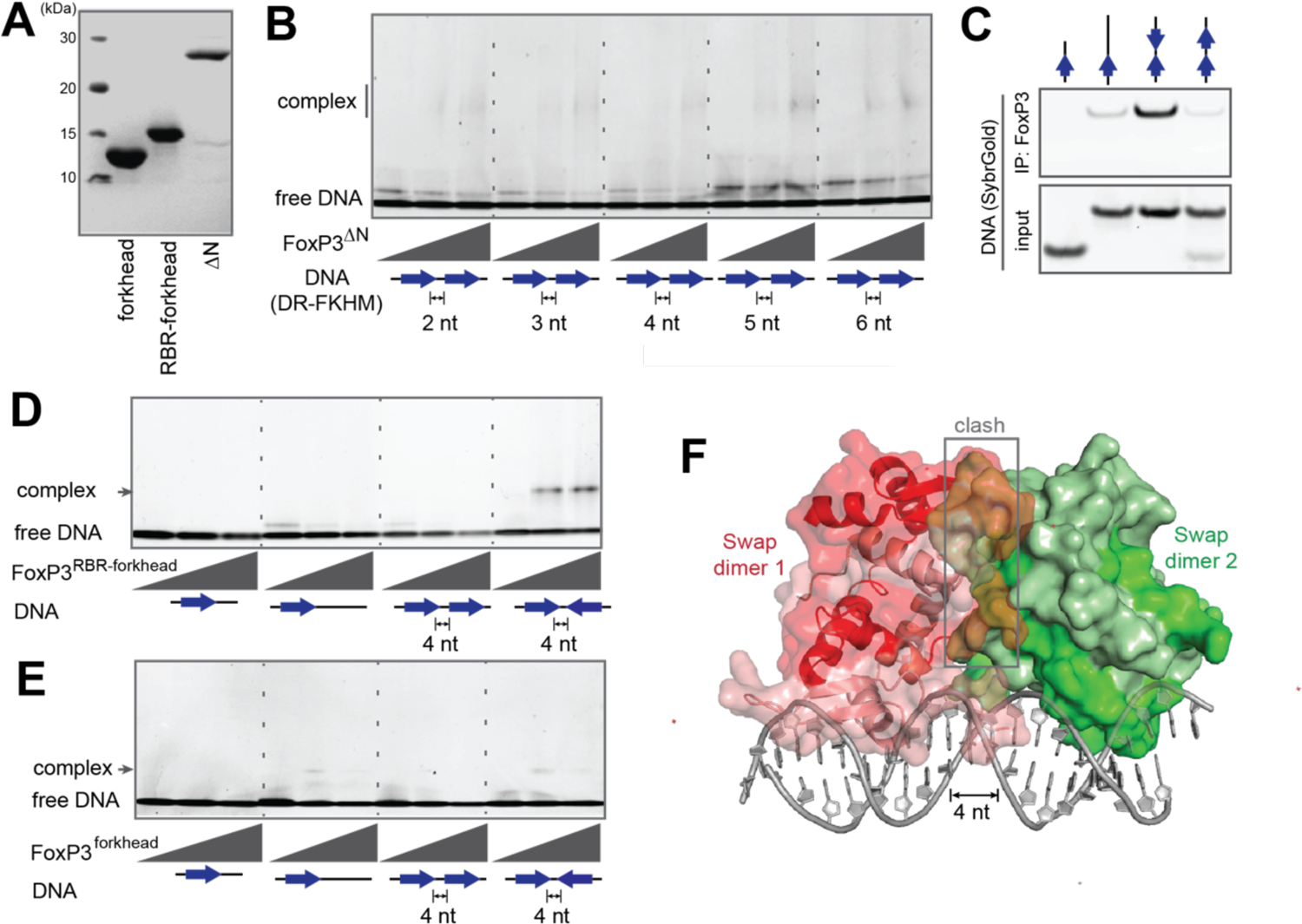
FoxP3^ΔN^ and FoxP3^RBR-forkhead^, but not FoxP3^forkhead^, preferentially binds IR-FKHM^4g^. Related to Figure 1. A. SDS-PAGE analysis of recombinant proteins FoxP3^ΔN^, FoxP3^RBR-forkhead^, and FoxP3^forkhead^ purified from *E. coli*. B. EMSA of FoxP3^ΔN^ (0, 0.4 and 0.8 μM) with DNA containing direct repeats (DR) of FKHM with various gap sizes (0.2 μM). Sybr Gold stain was used to visualize DNA. C. DNA specificity of full-length FoxP3 as measured by FoxP3 pull-down. HA-tagged FoxP3 was ectopically expressed in 293T cells and purified by anti-HA immunoprecipitation (IP). DNA was added to FoxP3 bound to beads and further purified to analyze FoxP3–DNA interaction. DNA in the eluate was analyzed by Sybr Gold stain. D. EMSA of FoxP3^RBR-forkhead^ (0, 0.4 and 0.8 μM) using four DNA oligos (0.2 μM) with different FKHM arrangements. E. EMSA of FoxP3^forkhead^ (0, 0.8 and 1.6 μM) using four DNA oligos (0.2 μM) with different FKHM arrangements. F. Modeling of the two swap dimers bound to IR-FKHM^4g^, where each dimer occupies a FKHM facing each other. Modeling suggests that the two swap dimers cannot simultaneously occupy the two FKHMs due to steric clash. Data in (B-E) are representative of at least three independent experiments.

**Figure S2.**
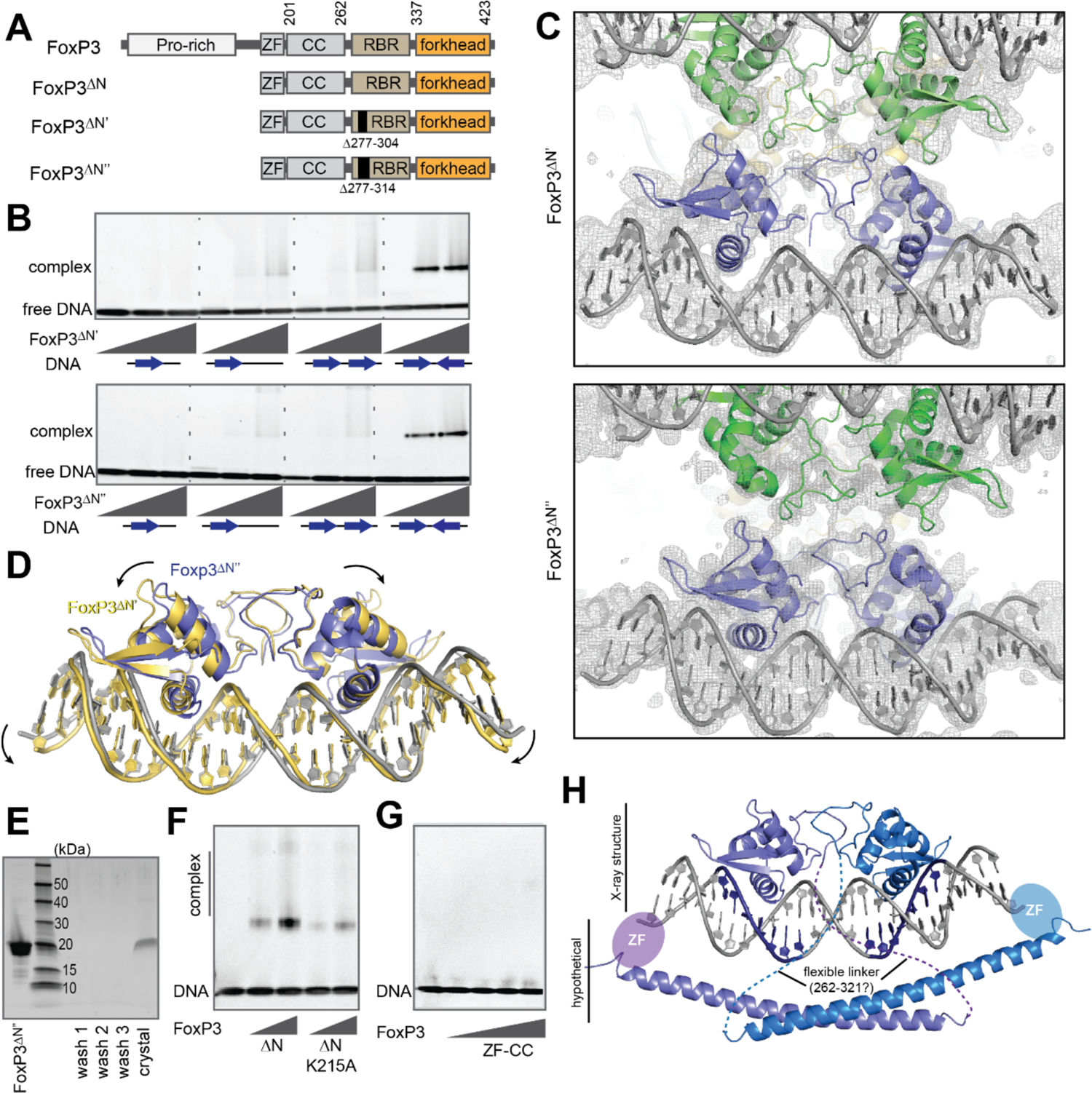
Crystal structures of FoxP3 in complex with IR-FKHM^4g^. Related to Figure 1. A. Crystallization constructs FoxP3^ΔN’^ and FoxP3^ΔN’’^, where residues 277-304 and 277-314 in the RBR linker, respectively, are deleted for crystallization. B. EMSA of FoxP3^ΔN’^ and FoxP3^ΔN’’^ (0, 0.4 and 0.8 μM) using four DNA oligos (0.2 μM) with different FKHM arrangements. FoxP3^ΔN’^ and FoxP3^ΔN’’^ also prefer IR-FKHM^4g^, as with FoxP3^ΔN^. C. Composite omit 2F_o_-F_c_ map for the crystal structures of FoxP3^ΔN’^ and FoxP3^ΔN’’^ (sigma=1). Purple proteins are one H-H dimer (bottom), while green (top) and yellow (back) proteins are two other H-H dimers related by the 3-fold crystallographic symmetry. Note that the two monomers within each H-H dimer were refined with crystallographic 2-fold symmetry using the reflection data processed with P6_3_22 symmetry. See Methods for details. D. Comparison of the FoxP3^ΔN’^ (yellow) and FoxP3^ΔN’’^ (purple) structures showing slightly altered interaction between monomers and DNA conformations. E. SDS-PAGE analysis of the crystal (FoxP3^ΔN’’^) showing that the protein is not proteolyzed. F. EMSA of FoxP3^ΔN^ (0, 0.4 and 0.8 μM) with and without K215A mutation in ZF. DNA with IR-FKHM^4g^ was used. K215 is a highly conserved Lys that in related C2H2-type ZF proteins is involved in DNA binding. G. EMSA of FoxP3^ZF-CC^ (0, 0.5, 1, 2 and 4 μM) using DNA with IR-FKHM^4g^. Note that DNA binding is barely detectable even with 4 μM of FoxP3^ZF-CC^. H. A model of how FoxP3 binds DNA. While ZF and CC could not be resolved in either crystal structure, our data suggest that ZF contributes to DNA binding and that ZF can bind nearby sites on DNA. The locations of ZF and CC relative to forkhead are likely heterogeneous, as evidenced by the lack of electron density for ZF and CC in either of our crystal structures. This is possible because ZF and CC are tethered to forkhead through a long RBR linker, the majority of which (residues 262-321) is also unresolved in the crystal structure and likely flexible. Note that a flexible, a 60-amino acid-long linker is long enough to wrap around DNA. Based on these data, we propose an overall architecture of FoxP3 where the H-H dimer of the forkhead domain forms a primary DNA binding unit, while two ZFs form a secondary binding unit flexibly tethered to the primary unit. Data in (B, F and G) are representative of at least three independent experiments.

**Figure S3.**
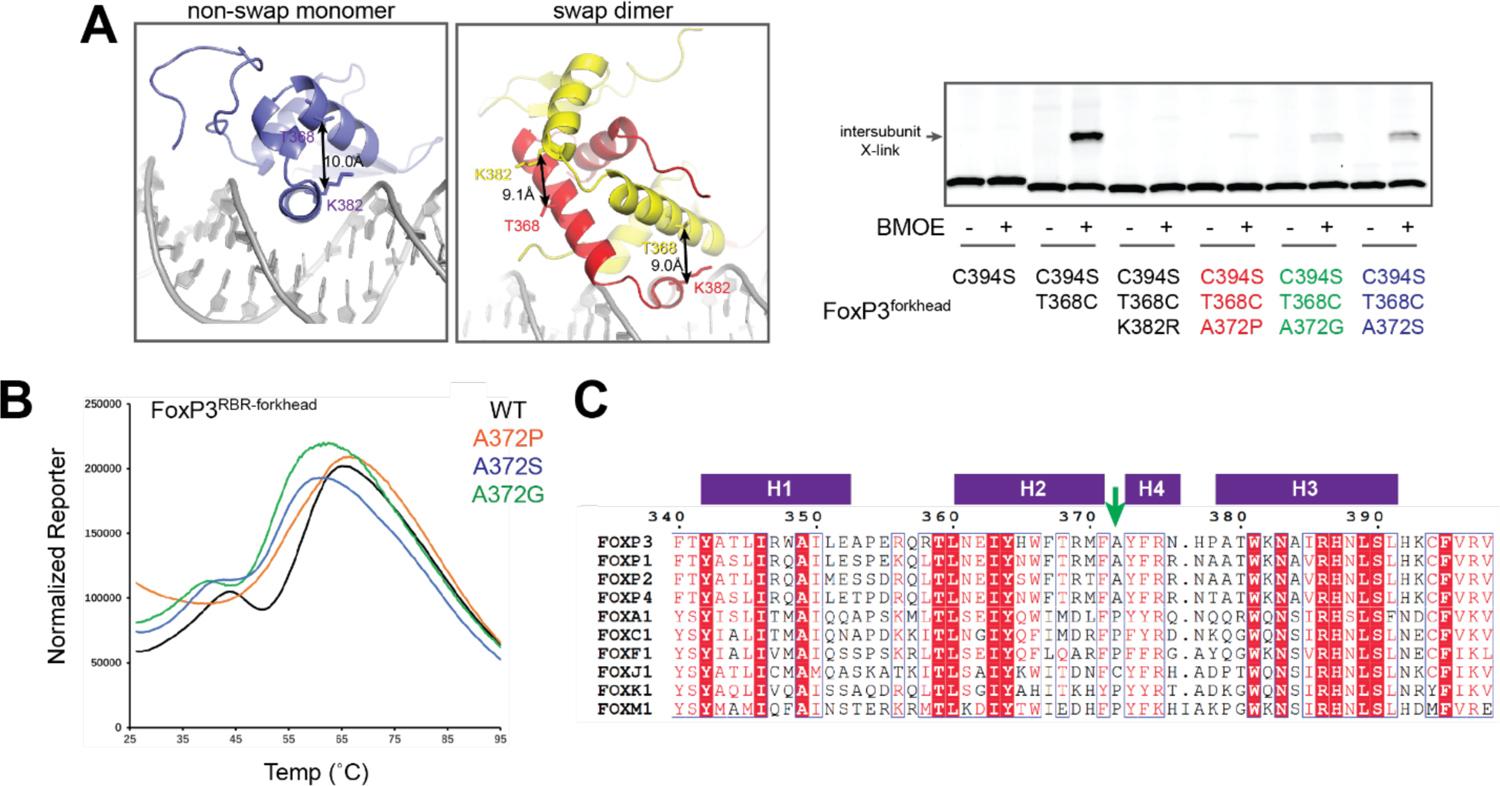
Effect of A372 mutations on swap dimerization and thermal stability. **Related to** Figure 2. A. Protein-protein crosslinking (X-linking) to examine the effect of A372 mutations on swap-dimerization. In order to design swap dimer-specific X-linking construct, we introduced C394S, which eliminates background X-linking by BMOE (bifunctional crosslinker that reacts with Cys or Lys). Additionally, we mutated T368 to Cys, which would lead to inter-subunit X-linking between T368C and K382 in the swap dimer, but intra-subunit X-linking in the non-swap monomer. Upon treatment with BMOE, we found that FoxP3^forkhead^ forms inter-subunit X-link in a manner that depends on T368C and K382 (right, lanes 1-6), consistent with its swap-dimeric structure. Introduction of A372P, A372G or A372S reduced the level of inter-subunit X-linking (right, lanes 7-12), suggesting that these mutations suppress swap dimerization. B. Thermal shift assay showing the reporter fluorescence level with increasing temperature. FoxP3^RBR-forkhead^ with and without mutations in A372 were compared. Even though A372 mutations do not alter the monomeric state of FoxP3^RBR-forkhead^ (Figure 2D), A372P shows a significantly altered melting curve compared to WT, A372G or A372S. This result suggests that A372P, but not A372G or A372S, have a negative impact on the thermal stability of FoxP3 independent of its ability to suppress swap dimerization. C. Sequence alignment of forkhead TFs showing conservation of Ala at the junction of H2 and H4 (green arrow, position 372 in FoxP3 numbering) within the FoxP family. Most forkhead TFs have Pro at the equivalent position although Cys was also observed for FoxJ1. Data in (A-B) are representative of at least three independent experiments.

**Figure S4.**
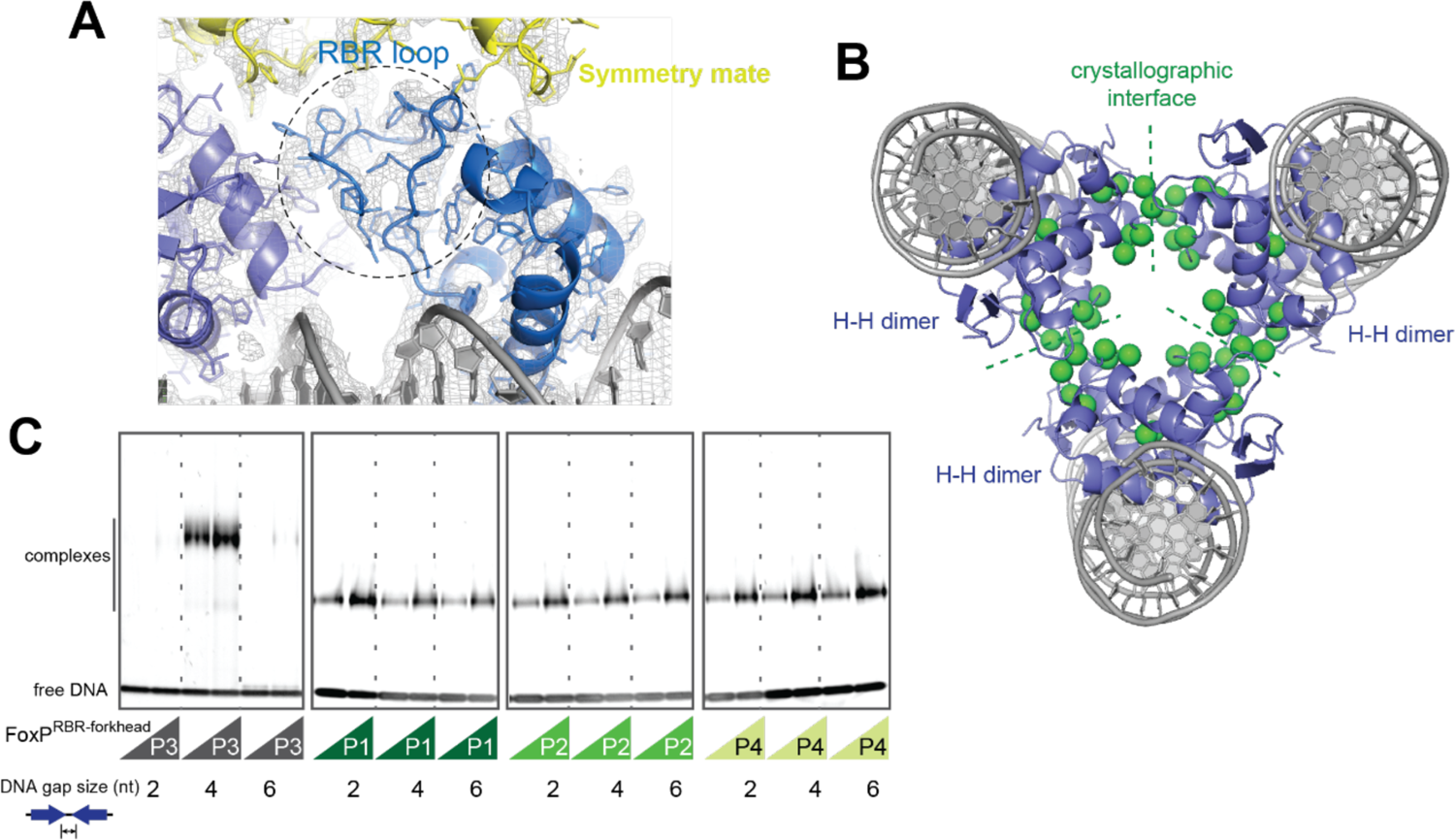
Crystallographic packing of FoxP3 and DNA specificity of FoxP1-4. Related to Figure 3. A. 2Fo-Fc map near the RBR loop. Symmetry mate related by the 3-fold rotation axis is shown in yellow. B. Crystal packing of FoxP3 looking down the 3-fold rotation axis. Hydrophobic residues in the RBR loop (indicated in Figure 3A) are shown in green spheres. C. EMSA of NusA-tagged FoxP1-4 (0.4 and 0.8 μM) using DNA oligos (0.2 μM) harboring IR-FKHM with a 2, 4 and 6 nt gap. All proteins were RBR-forkhead domains fused with NusA. Data in (C) are representative of at least three independent experiments.

**Figure S5.**
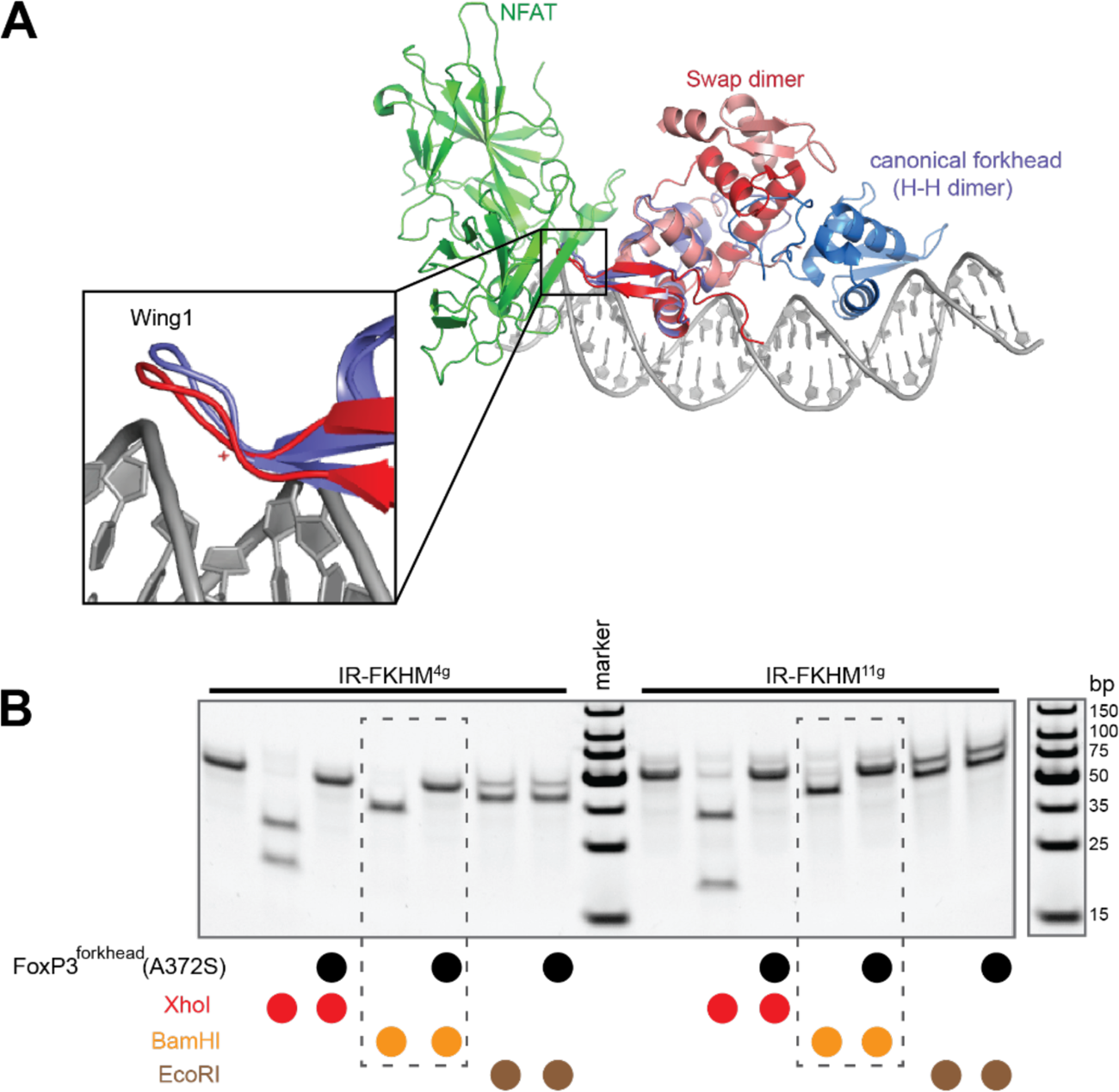
FoxP3–DNA interaction in the presence of NFAT. Related to Figure 4. A. Overlay of the swap dimeric and non-swap monomeric FoxP3, showing that the NFAT interface of FoxP3 (wing 1) is nearly identical in both structures. Note that the structure of NFAT (green) was determined in complex with the swap dimer (PDB: 3QRF). B. Restriction enzyme protection assay with FoxP3^forkhead^ (6.4 μM) in the presence of NFAT (0.4 μM). The mutant A372S was used to ensure that forkhead forms the non-swap conformation. Experiments were performed as in Figure 4D. The result shows that FoxP3^forkhead^ (A372S) binds IR-FKHM^11g^ with individual forkhead occupying each FKHM (as in model (i) in Figure 4C). This differs from FoxP3^ΔN^, which binds IR-FKHM^11g^ as a H-H dimer (as in model (ii) in Figure 4C). Data in (B) are representative of at least three independent experiments.

**Figure S6.**
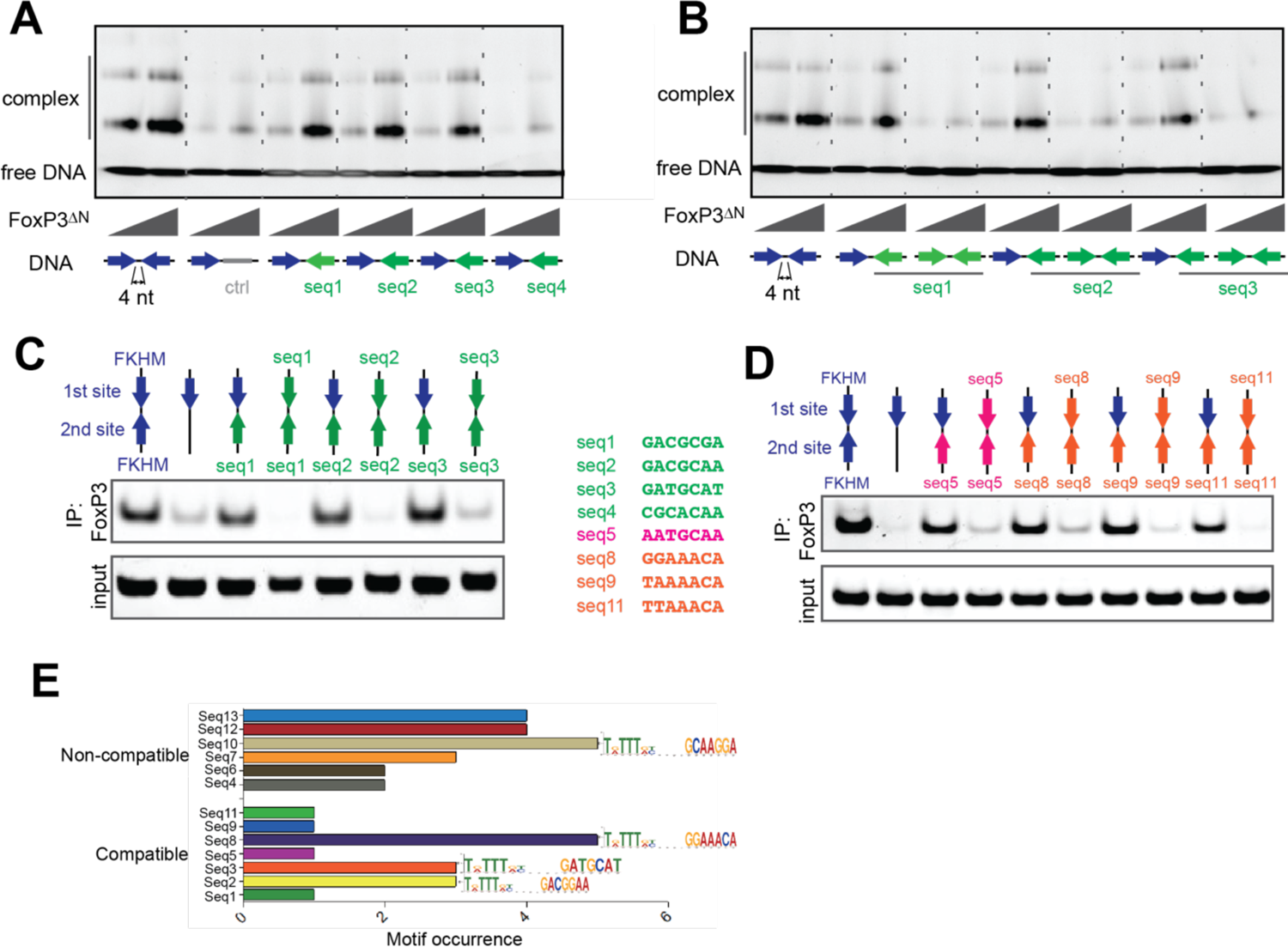
Head-to-head dimerization enables FoxP3 to recognize diverse sequences. Related to Figure 5. A. EMSA of FoxP3^ΔN^ (0.4 and 0.8 μM) using DNA containing IR-FKHM^4g^ or FKHM paired with seq1-4 (from Figure 5C). B. EMSA of FoxP3^ΔN^ (0.4 and 0.8 μM) using DNA containing IR-FKHM^4g^ or seq1-3 in inverted repeat with 4 nt gaps. C. FoxP3 interaction with DNA harboring seq1-3 paired with FKHM or inverted repeats of seq1-3. 4 nt gap was used for all. D. FoxP3 interaction with DNA harboring seq5/8/9/11 (from Figure 5D) in inverted repeats *vs.* those paired with FKHM. E. Occurrence of compatible and non-compatible dimers in FoxP3 ChIP-seq binding sites that include an FKHM (n=548). Motif occurrences were counted with a FIMO p-value<10^-5^ and score greater than 12. Data in (A-D) are representative of at least three independent experiments.

**Figure S7.**
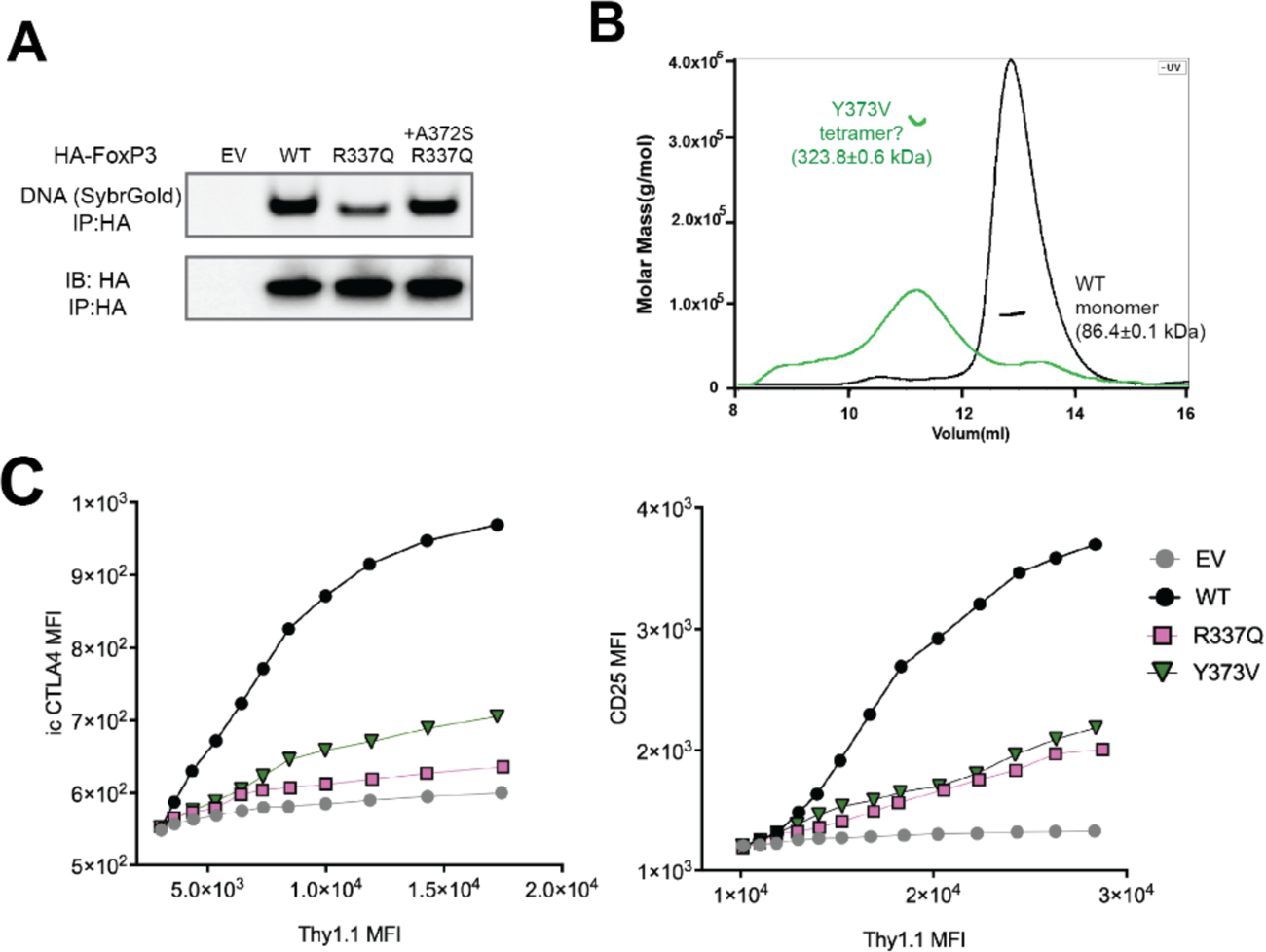
Analyses of IPEX mutations R337Q and Y373V. A. FoxP3 interaction with DNA. HA-tagged, full-length FoxP3 was ectopically expressed in 293T cells and purified by anti-HA immunoprecipitation (IP). Equal amount of IR-FKHM^4g^ DNA was added to FoxP3 bound to beads and further purified to analyze FoxP3–DNA interaction. Bound DNA and proteins were isolated and analyzed by native-PAGE and SDS-PAGE, respectively. Sybr Gold stain was used for DNA measurement. B. SEC-MALS of NusA-tagged FoxP3^RBR-forkhead^ (WT and Y373V). C. Transcriptional activity of FoxP3 with R337Q or Y373V. Experiments were performed as in Figure 2C.

**Table S1.**
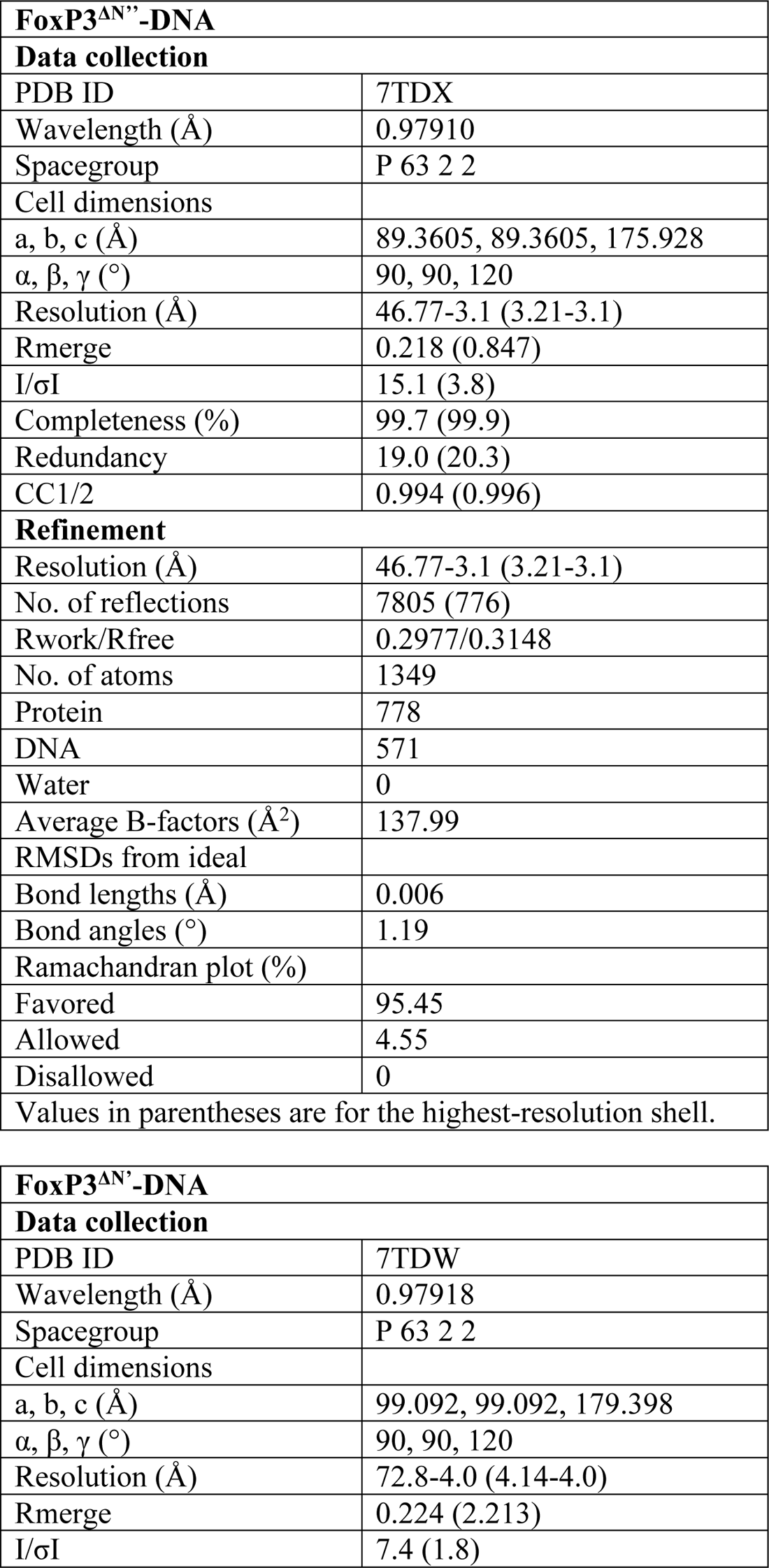
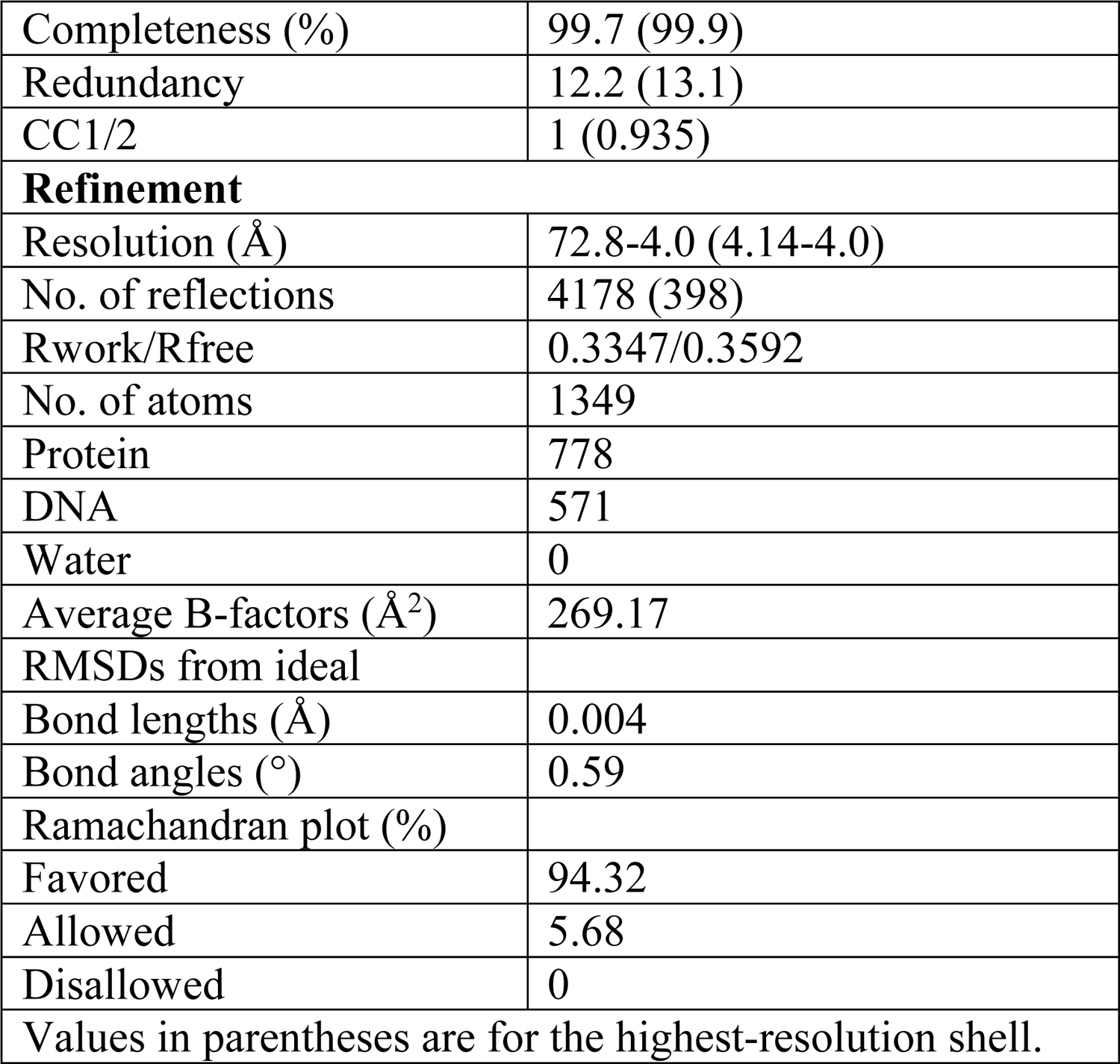
Data collection and refinement statistics

**Table S2.**
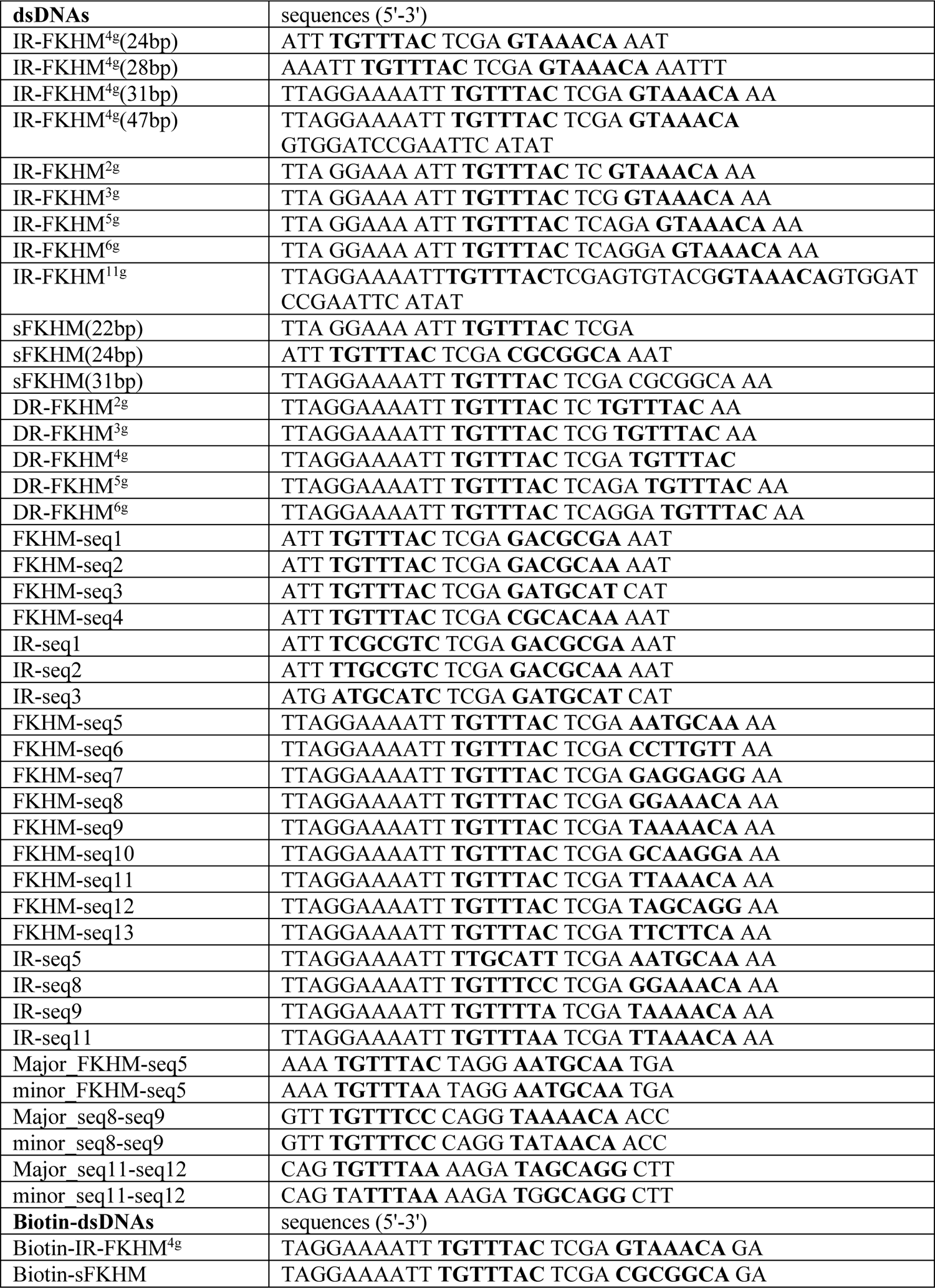
Sequences of dsDNAs and Biotin-dsDNAs. Related to STAR methods.

## KEY RESOURCES TABLE

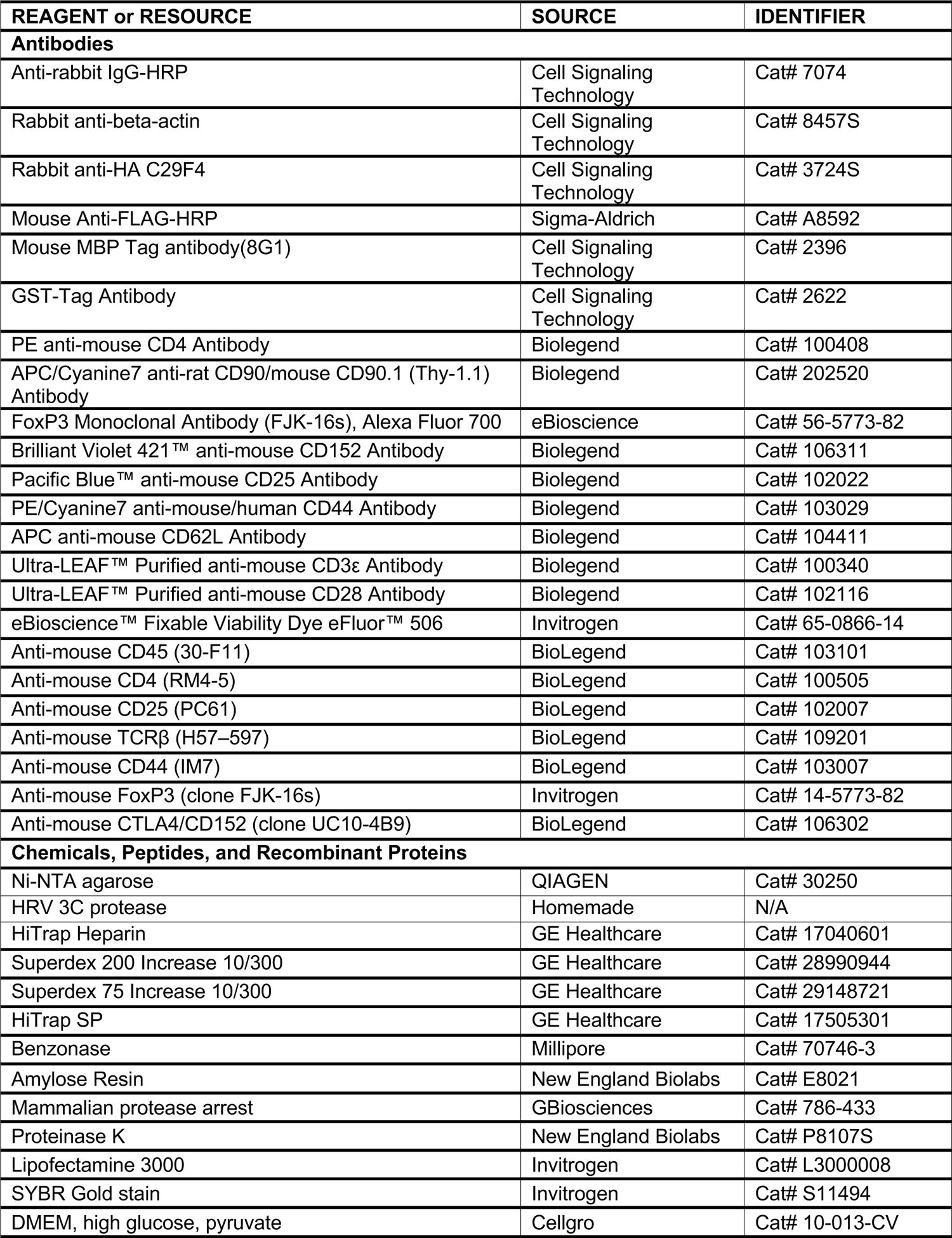
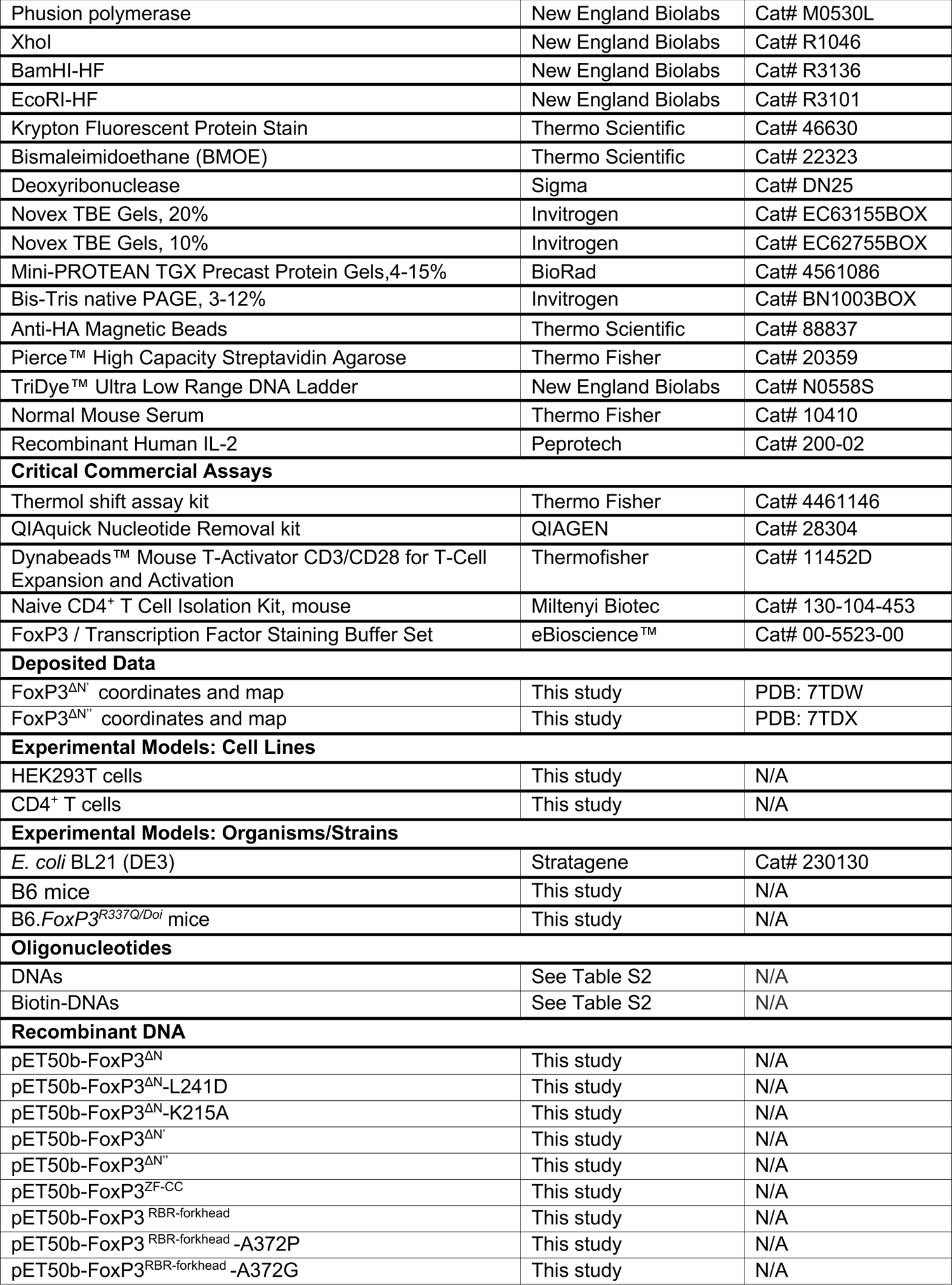
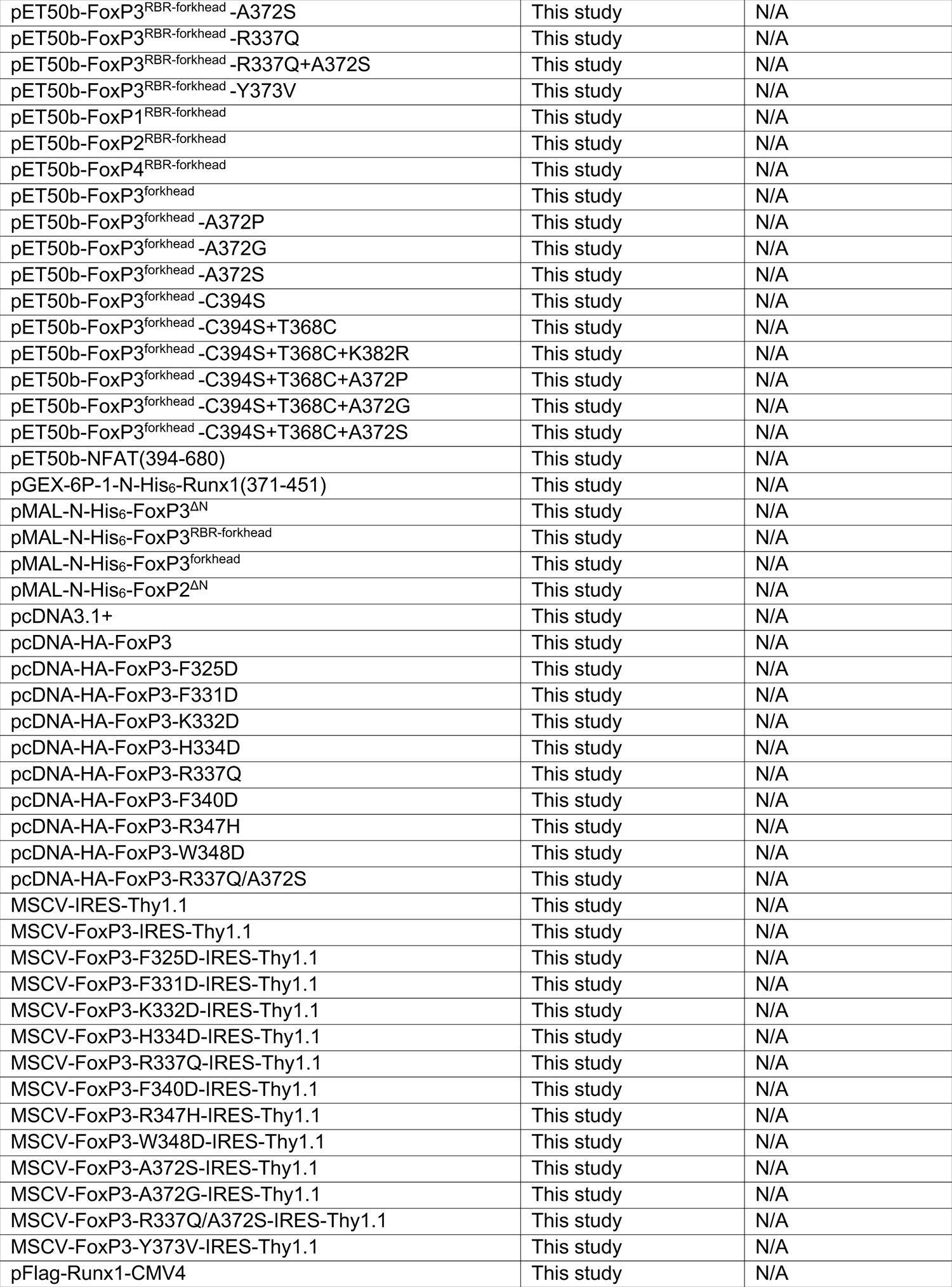
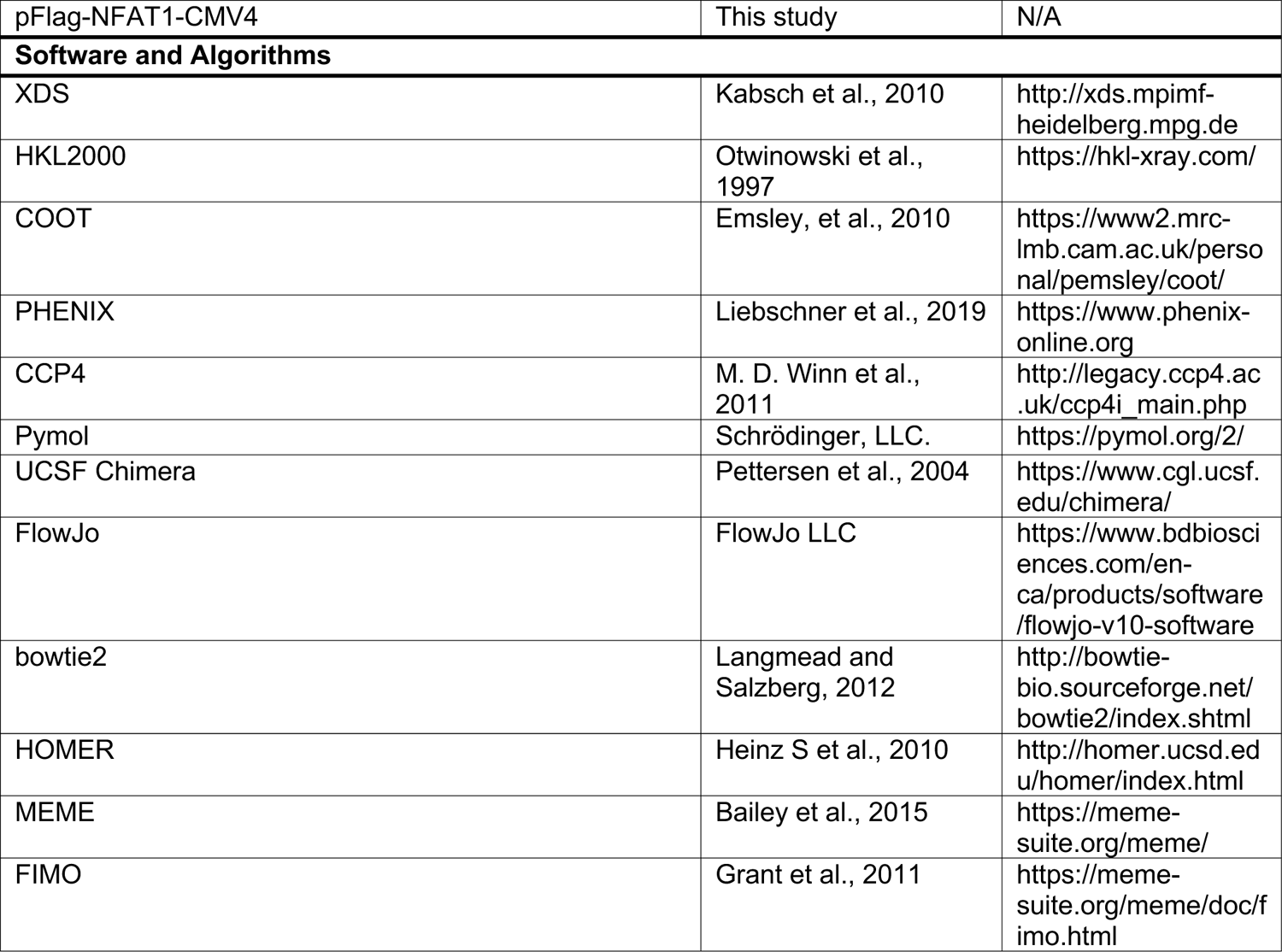

### RESOURCE AVAILABILITY

#### Lead contact

Further information and requests for resources and reagents should be directed to and will be fulfilled by the Lead Contact, Sun Hur

### Materials Availability

All plasmids generated in this study are available from the Lead Contact with a completed Materials Transfer Agreement.

### Data and Code Availability

The atomic coordinates have been deposited in the Protein Data Bank with accession codes: 7TDW and 7TDX.

## EXPERIMENTAL MODEL AND SUBJECT DETAILS

### HEK293T cells

Cells were maintained in DMEM (High glucose, L-glutamine, Pyruvate) with 10% fetal bovine serum, 1% penicillin/streptomycin.

### Naive CD4^+^ T Cell**s**

Cells were isolated by using Naive CD4^+^ T Cell Isolation Kit (Miltenyi Biotec, Cat#130-104-453) according to the manufacturer’s instructions and maintained in complete RPMI medium (10% FBS heat-inactivated, 2mM L-Glutamine, 1mM Sodium Pyruvate, 100μM NEAA, 5mM HEPES, 0.05mM 2-ME).

### Mice

B6.*FoxP3^R337Q/Doi^* mice were generated by CRISPR mutagenesis as described elsewhere (Leon et al, in preparation). Briefly, the *Foxp3* allele carries two consecutive point-mutations of guanine to adenine and adenine to guanine at position 1286-1287 in exon 11 (NM_054039.2), resulting in substitution of arginine for glutamine at amino acid 337 (R337Q), first amino-acid of the DNA-binding forkhead domain (Figure 1A). The germline mutation was maintained onto the B6 background and bred in the SPF facility at Harvard Medical School. For the experiments reported here, 8-week-old R337Q males and their WT B6 littermates were analyzed. Data from two independent litters were pooled. All experiments were performed following animal protocols approved by the HMS Institutional Animal Use and Care Committee (protocols IS00000054).

## METHOD DETAILS

### Material Preparation

#### Plasmids

Mouse FoxP3 was used for all analyses. For bacterial expression of FoxP3 variants, the genes encoding mouse FoxP3^ΔN^ (residues 188-423), FoxP3^RBR-forkhead^ (residues 284-423), FoxP3^forkhead^ (residues 336-423), FoxP3^ZF-CC^ (residues 188-284) were inserted into pET50b between Xmal and HindIII sites, and into a modified pMAL-c2 vector between BamHI and Xbal sites respectively. All mutations within FoxP3 were generated by site-directed mutagenesis using Phusion High Fidelity (New England Biolabs) DNA polymerases. For crystallization, the gene encoding mouse FoxP3^ΔN’^ (residues 204-276+305-417) and FoxP3^ΔN”^ (residues 204-276+315-417) were cloned into pET50b using overlap extension PCR. Mouse FoxP1^RBR-forkhead^ (residues 433-579), mouse FoxP2^ΔN^ (residues 353-588), mouse FoxP2^RBR-forkhead^ (residues 447-588) and mouse FoxP4^RBR-forkhead^ (residues 418-577) were cloned into pET50b between Xmal and HindIII sites, and modified pMAL-c2 vector between BamHI and Xbal sites respectively. Mouse Runx1 (residues 371-451) was cloned into pGEX-6P-1 between EcoRI and XhoI. Mouse NFAT1 (residues 394-680) was cloned into pET50b plasmid between Xmal and HindIII sites.

For Mammalian expression plasmids, HA-tagged mouse FoxP3 CDS was inserted into pcDNA3.1+ vector between KpnI and BamHI sites. All FoxP3 mutations including F325D, F331D, K332D, H334D, R337Q, R347H, W348D and R337Q/A372S were generated by site-directed mutagenesis using Phusion High Fidelity (New England Biolabs) DNA polymerases. The genes encoding mouse Runx1 and NFAT1 were inserted into pFlag-CMV4 vector between EcoRI and XbaI, and between NotI/XbaI sites respectively. For retroviral packaging plasmids, HA-tagged mouse FOXP3 CDS was inserted into MSCV-IRES-Thy1.1 vector. All FoxP3 mutations including F325D, F331D, K332D, H334D, R337Q, R347H, W348D, A372S, A372G, R337Q/A372S and Y373V were generated by site-directed mutagenesis using Phusion High Fidelity (New England Biolabs) DNA polymerases.

#### DNAs

Single-stranded DNA oligos were synthesized by IDTDNA. Double-stranded DNAs for EMSA assay, pulldown assay and restriction enzyme protection assay were annealed from single-stranded, complementary oligos. After briefly spinning down each oligonucleotide pellet, ssDNAs were dissolved in annealing buffer (10 mM Tris pH 7.5, 50 mM NaCl). Complementary ssDNAs were then mixed together in equal molar amounts, heated to 94°C for 2 minutes and gradually cool down to room temperature. For dsDNAs used in crystallization, HPLC purified single-stranded, complementary oligos were purchased from IDTDNA. After annealing, dsDNA was further purified by size-exclusion chromatography on Superdex 75 Increase 10/300 (GE Healthcare) columns in 20 mM Tris pH 7.5, 150 mM NaCl. Biotin labeled ssDNA were synthesized by IDTDNA and then dissolved in annealing buffer (10 mM Tris pH 7.5, 50 mM NaCl). Complementary biotin labeled ssDNAs were then mixed together in equal molar amounts, heated to 94°C for 2 minutes and gradually cool down to room temperature.

#### Protein expression and purification

All recombinant proteins in this paper were expressed in BL21(DE3) at 18°C for 16-20 hr following induction with 0.2 mM IPTG. Cells were lysed by high-pressure homogenization using an Emulsiflex C3 (Avestin). All protiens are from the *Mus. musculus* sequence. FoxP3^ΔN^, FoxP3^ΔN’^ and FoxP3^ΔN”^ were expressed as a fusion protein with N-terminal His_6_-NusA tag. After purification using Ni-NTA agarose, the proteins were treated with HRV3C protease to cleave the His_6_-NusA-tag and were further purified by a combination of chromatography using HiTrap Heparin (GE Healthcare), Hitrip SP (GE Healthcare) and Superdex 200 Increase 10/300 (GE Healthcare) columns. FoxP3^RBR-forkhead^, FoxP3^forkhead^ and NFAT1 (residues 394-680) were expressed as fusion protein with N-terminal His_6_-NusA tag. After purification using Ni-NTA agarose, the proteins were treated with HRV3C protease to cleave the His_6_-NusA-tag and were further purified by size-exclusion chromatography on Superdex 75 Increase 10/300 (GE Healthcare) column. His_6_-MBP fused FoxP3^ΔN^, FoxP3 ^RBR-forkhead^, FoxP2^ΔN^ and His_6_-GST fused Runx1 (residues 371-451) were purified by a combination of chromatography on Ni-NTA agarose and Superdex 200 Increase 10/300 (GE Healthcare) columns. For SEC-MALS analysis, proteins were purified without HRV3C cleavage to increase the molecular weight of the protein for accurate size determination. His_6_-NusA fused FoxP3 variants, FoxP1^RBR-forkhead^, FoxP2^RBR-forkhead^ and FoxP4^RBR-forkhead^ were purified by Ni-NTA agarose affinity chromatography, followed by size-exclusion chromatography on Superdex 200 Increase 10/300 (GE Healthcare) columns.

#### Electrophoretic Mobility Shift Assay (EMSA)

0.2 μM of DNA was mixed with the indicated amount of FoxP3 variants or other FoxP proteins in the buffer (20 mM Tris, pH 7.5, 150 mM NaCl, 1.5 mM MgCl_2_, and 2 mM DTT). Proteins and DNAs were incubated for 30 min at 4 °C and analyzed on 3-12% gradient Bis-Tris native gels (Life Technologies) at 4 °C. After staining with Sybr Gold stain (Life Technologies), Sybr Gold fluorescence was recorded using iBright FL1000 (Invitrogen) and analyzed with iBright Analysis Software.

#### Multi-Angle Light Scattering (MALS)

NusA-fused FoxP3 variants, FoxP1^RBR-forkhead^, FoxP2^RBR-forkhead^ and FoxP4^RBR-forkhead^ were analyzed with Superdex 200 Increase 10/300 column (GE Healthcare), which was connected to a miniDAWN MALS detector (Wyatt Technology) and an Optilab differential refractive index (dRI) detector (Wyatt Technology). The buffer (20 mM Tris pH7.5, 500 mM Nacl, 2 mM DTT) was used for size-exclusion chromatography. The AKTA pure (GE Healthcare)’s UV 280 nm absorbance signal was used for concentration detection. The Optilab differential refractive index (dRI) detector measured dRI data for additional concentration analysis. Data analysis and MW calculations were performed using the ASTRA7.3.1 software (Wyatt Technology).

#### Crystallization of FoxP3^ΔN’^ and FoxP3^ΔN”^

Crystallization of FoxP3 was tried with various constructs and DNA, but crystals were only obtained with FoxP3^ΔN’^ and FoxP3^ΔN”^, which had C217S and C231S mutations and internal deletions in the RBR loop. FoxP3^ΔN’^ contained residues 204-276 and 305-417, while FoxP3^ΔN”^ contained residues 204-276 and 315-417. Purified proteins (post Superdex 200 Increase) were mixed with purified IR-FKHM^4g^ dsDNA (5’-AAATT<u>TGTTTAC</u>TCGA<u>GTAAACA</u>AATTT, post Superdex 75 Increase 10/300) at 1:1.2 molar ratio in 20 mM Tris-Hcl pH 7.5, 150 mM Nacl, 3 mM 2-mercaptoethanol and the mixture was concentrated to 10 mg/ml (Protein concentration) using an Amicon Ultra-4 filter (3 kDa molecular-weight cutoff, Millipore). The FoxP3^ΔN’^–DNA complex was then mixed with the reservoir solution (0.1 M Tris-Hcl pH 8.5, 12% PEG4000) at a 1:1 volume ratio and was crystallized at 18°C by vapor diffusion using the hanging drop method. Crystals for the FoxP3^ΔN”^–DNA complex grew in the reservoir solution (0.5 M Lithium sulfate, 2% PEG8000) and were obtained using a similar method as with FoxP3^ΔN’^. To test whether the FoxP3^ΔN’’^ protein was intact in the crystal, a single crystal was picked, washed three times in 10 μl reservoir solution and then directly transferred into 10 μl of 1x SDS sample buffer prior to SDS-PAGE analysis. The protein was visualized by Krypton stain (Thermo Scientific).

#### Data Collection, Structure Determination, and Analysis

Crystals were cryoprotected in the reservoir solution supplemented with 25% glycerol and were flash-cooled in liquid nitrogen. The X-ray diffraction data were collected at the Advance Photon Source, beamline NECAT-24-ID-E. Diffraction data were processed using XDS (Kabsch, 2010) and HKL2000 (Otwinowski and Minor, 1997) in P6_3_22 symmetry. The FoxP3^ΔN”^–DNA complex structure was solved by molecular replacement with Phaser (McCoy et al., 2007) using canonical forkhead structures of FoxN1 (PDB 6EL8) and ∼14 bp DNA molecule (from PDB: 3QRF). The known structure of FoxP3 forkhead domain (PDB: 3QRF) and FoxP3 coiled-coil domain (PDB: 4I1L) were also tried as molecular replacement templates, but no solution was obtained. The MR solution for DNA showed that two 14 bp DNAs face each other through 2-fold symmetry in a way that can make one continuous 28 bp DNA, which was the biological sample used for crystallization. Note that the data processed with P6_3_ symmetry showed a single FoxP3 dimer bound to 28 bp DNA in the asymmetric unit, of which the structure was nearly identical to that obtained with P6_3_22. However, data merging statistics and quality of the electron density map were inferior to those obtained with P6_3_22. Therefore, P6_3_22 was used for subsequent model building and refinment. The model of FoxP3^ΔN”^monomer with 14bp DNA was automatically built using Autobuild (Terwilliger et al., 2008), followed by manual model building using COOT (Emsley et al., 2010). Structural refinement was performed with PHENIX (Liebschner et al., 2019) and REFMAC5 in ccp4 (Murshudov et al., 2011). The FoxP3^ΔN’^–DNA complex structure was solved by molecular replacement using the model of the FoxP3^ΔN”^–DNA complex, followed by structural refinement using PHENIX and REFMAC5. The atomic coordinates of FoxP3^ΔN’^– DNA and FoxP3^ΔN”^–DNA have been deposited in the Protein Data Bank with accession codes 7TDW and 7TDX, respectively. Data collection and refinement statistics are summarized in Table S1. Figures of structure illustration were prepared using Pymol (Schrödinger, LLC).

#### Co-IP of HA-FoxP3 and FLAG-Runx1

HEK293T cells (in 6-well plate) were transfected with empty vector or pcDNA encoding HA-tagged FoxP3 (wild-type or mutants), or pFlag-Runx1-CMV4 respectively using Lipofectamine 3000 (Life Technologies) following manufacturer’s instructions. 48 hours later, cells were washed twice in 1XPBS (1 ml of PBS for each well) and spun down at 500g for 5 mins. Cells were resuspended in 500 μl lysis buffer (20 mM HEPES pH 7.5, 0.05% IGEPAL, 1.5 mM MgCl_2_, 10 mM KCl, 5 mM EDTA and 1x Mammalian protease cocktail) for 15 minutes at 4 °C and were spun down at 500g for 5 minutes at 4 °C. The supernatant was collected and centrifuged at 14000 rpm for another 5 minutes. Soluble fractions containing HA-FoxP3 and Flag-Runx1 (400 μl each) were mixed together and incubated with anti-HA magnetic beads (4 μl) (Thermo Scientific) for 1 hour at 4°C with slow rotation. To examine if FoxP3–Runx1 interaction was dependent on nucleic acids, 4 μl or 8 μl of Benzonase (Millipore) was added to the FoxP3 and Runx1 mixture for 10 minutes at 25 °C. Beads were washed three times with RIPA buffer (50 mM Tris-HCl pH 7.5, 1 mM EDTA, 1% Triton x-100, 0.5% Sodium Deoxycholate, 0.1% SDS, 150 mM NaCl, 1x Mammalian protease cocktail), followed by protein elution using SDS loading buffer and analysis by SDS-PAGE.

#### Co-IP of FoxP3 and DNAs

HEK293T cells were transfected with pcDNA encoding HA-tagged FoxP3 (wild-type or mutants). After 48 hours, cells were lysed using RIPA buffer (10mM Tris-HCl, pH 8.0, 1mM EDTA, 1% Triton X-100, 0.1% Sodium Deoxycholate, 0.1% SDS, 140 mM NaCl and 1x proteinase inhibitor) and treated with Benzonase (Millipore) for 30 mins. The lysate was then incubated with Anti-HA Magnetic Beads (Thermo Fisher) for 1 hour. Beads were washed three times using RIPA buffer and incubated with DNA oligos for 20 mins at room temperature. Bound DNA was recovered using proteinase K (New England Biolabs), purified using QIAquick Nucleotide Removal kit (QIAGEN) and analyzed on 10% Novex TBE Gels (Invitrogen).

#### MBP-FoxP3^ΔN^ pulldown with DNAs

Purified MBP-mFoxP3^ΔN^ or MBP-FoxP2^ΔN^ protein (0.4 μM) was incubated with DNA (0.1 μM) in the buffer (20 mM Tirs, pH 7.5, 100 mM NaCl, 1.5 mM MgCl_2_) for 20 mins at RT, and was further incubated with Amylose Resin (25 μL) (New England Biolabs) with rotation for 30 mins at RT. Bound DNA was recovered using proteinase K (New England Biolabs), purified using QIAquick Nucleotide Removal kit (QIAGEN) and analyzed on 10% Novex TBE Gels (Invitrogen).

#### Biotin-DNAs pulldown with HA-FoxP3 variants

HEK293T cells were transfected with pcDNA-HA-FoxP3 (wild-type or mutations) and lysed using RIPA buffer (10 mM Tris-HCl, pH 8.0, 1 mM EDTA, 1% Triton X-100, 0.1% Sodium Deoxycholate, 0.1% SDS, 140 mM NaCl and 1x proteinase inhibitor). Biotin-dsDNA (1 μM) was incubated with the lysate for 1 hour. Streptavidin Agarose (25 μL)(Thermo Fisher) was added and further incubated for 30 mins with rotation. Beads were washed three times using RIPA buffer and eluted using SDS loading buffer prior to SDS-PAGE analysis.

#### MBP-FoxP3^RBR-forkhead^ pulldown with GST-Runx1(371-451)

Purified MBP-FoxP3^RBR-forkhead^ protein (1 μM) was incubated with purified GST-Runx1(371-451) protein (0.5 μM) with or without 2 μM DNA for 30 mins at room temperature in RIPA buffer (10 mM Tris-HCl, pH 8.0, 1 mM EDTA, 1% Triton X-100, 0.1% Sodium Deoxycholate, 0.1% SDS, 140 mM NaCl and 1x proteinase inhibitor). Streptavidin Agarose beas (25 μL) (Thermo Fisher) were added and incubated with rotation for 30 mins at RT. Beads were washed three times with RIPA buffer, prior to elution and analysis by SDS-PAGE or TBE gels.

#### Crosslinking Analysis

Protein-protein crosslinking using BMOE (Thermo Scientific) was carried out according to the product manual. Briefly, BMOE was added to 2 μM of FoxP3^forkhead^ protein to a final concentration of 100 μM in 1XPBS. After 1-hour incubation at 25°C, DTT (10 mM) was added to quench the crosslinking reaction. Samples were then analyzed by SDS-PAGE and Krypton staining (Thermo Scientific).

#### Thermal shift assay

FoxP3^RBR-forkhead^ (19 μl, 0.25 mg/ml) was mixed with 20x Protein thermal shift dye (1 μl) (Thermo Scientific) in SEC buffer (50 mM Tris-HCl pH 7.5, 150 mM Nacl, 2 mM DTT) and aliquoted into MicroAmp Optical 96-Well Reaction Plate (Applied Biosystems). Thermal shift assay was performed using StepOnePlus Real-Time PCR System (Applied Biosystems). Briefly, temperature was increased in a step-and-hold manner from 25°C to 98°C in a 0.3°C/cycle increment and with an equilibration time of 15s at each temperature. Normalized reporter (Rn) view visualizing the rise in fluorescence throughout the temperature ramp were generated by onestep plus software. The normalized reporter (Rn), displayed on the y-axis, is calculated as the fluorescence signal from the reporter dye normalized to the fluorescence signal of the passive reference.

#### Restriction enzyme protection assay

IR-FKHM^4g^ and IR-FKHM^11g^ (0.2 μM) were pre-incubated with NFAT (0.4 μM) for 10 mins at 25°C in the buffer (50 mM Potassium Acetate, 20 mM Tris-acetate, 10 mM Magnesium Acetate, 100 ug/ml BSA, pH 7.9, 50 mM NaCl). Then FoxP3^ΔN^ (1.6 μM for IR-FKHM^4g^ and 3.2 μM for IR-FKHM^11g^) was added to the mixture. Different concentration of FoxP3 was used for the two DNA because IR-FKHM^11g^ requires higher amount of FoxP3 for binding. After 10 mins of incubation at 25°C, restriction enzymes (XhoI, BamHI-HF or EcoRI-HF, New England Biolabs) were added and further incubated for 5 mins at 37°C. Restriction digestion was quenched by adding 1 μl proteinase K (New England Biolabs) and incubation for 2 mins at 37°C, followed by adding 0.25% SDS and 20 mM EDTA. DNA was analyzed on 20% Novex TBE Gels (Invitrogen). For restriction enzyme protection assay with FoxP3^forkhead^-A372S, 6.4 μM of FoxP3^forkhead^-A372S was used for both IR-FKHM^4g^ and for IR-FKHM^11g^.

#### CD4^+^ T Cell Cultures and Retroviral Transductions

Naïve CD4^+^ T cells were isolated by negative selection from mouse spleens by using the isolation kit (Miltenyi Biotec) according to the manufacturer’s instructions. Cell purity was validated with >90% by FACS analysis using PE anti-CD4 (Biolegend). Cells were then activated with anti-CD3 (Biolegend), anti-CD28 (Biolegend) and 50 U/mL of IL2 (Peprotech) in complete RPMI medium (10% FBS heat-inactivated, 2 mM L-Glutamine, 1 mM Sodium Pyruvate, 100 μM NEAA, 5 mM HEPES, 0.05 mM 2-ME). Stimulation of naïve CD4^+^ T cells was confirmed by their increased cell sizes and expression of the activation marker CD44 (BioLegend) by FACS analysis. After 48 hours, cells were spin-infected with retrovirus containing supernatant from HEK293T cells transfected with retroviral expression plasmids (Empty MSCV-IRES-Thy1.1 vector, wildtype-FoxP3 and mutations encoding vectors) and cultured for 2∼3 days in complete RPMI medium with 100 U/mL of IL2.

#### Flow Cytometry

For detecting of CD25 and CTLA4 expression in transduced CD4^+^ T cells, activated CD4^+^ T cells were stained with cell surface anti-CD25 (Biolegend) and Thy1.1 (Biolegend) on day 2 post retroviral infection. For intracellular staining, anti-CTLA4 (Biolegend) was applied on day 3 post retroviral infection using the Transcription Factor Staining Buffer Set (eBioscience) according to the manufacturer’s instructions. Flow cytometry data were analyzed with FlowJo software and presented as plots of mean fluorescence intensity (MFI) of CD25 and CTLA4 in cells grouped into bins of Thy1.1 intensity. Each result is representative of 3 independent experiments.

For analysis of mutant mice, single cell suspensions were obtained from murine spleens after physical dissociation with a 40 μm mesh and red blood cell lysis, and from the colonic lamina propria per (Sefik et al., 2015). After Fc blocking, extracellular staining was done in ice-cold buffer (phenol red–free DMEM, 2% FBS) for 30 min using antibodies against CD45 (30-F11; BioLegend, dilution 1:200), CD4 (RM4-5; BioLegend, dilution 1:200), CD25 (PC61; BioLegend, 1:50), TCRβ (H57–597; BioLegend, 1:150), CD44 (IM7; BioLegend, dilution 1:100). Cells were then fixed overnight at 4°C using 100 μL of Fix/Perm buffer (eBioscience), followed by permeabilization using 1X permeabilization buffer (eBioscience) for 45 minutes at room temperature in the presence of the following intracellular antibodies: FoxP3 (clone FJK-16s, Invitrogen, 1:75); CTLA4/CD152 (clone UC10-4B9, BioLegend, dilution 1:200). Data was recorded on a FACSymphonyTM flow cytometer (BD Biosciences) and analyzed using FlowJo 10 software.

#### FoxP3 ChIP-seq analysis

FoxP3 ChIP-seq (Kitagawa et al., 2017; Samstein et al., 2012) data was mapped to mm10 using bowtie2 (Langmead and Salzberg, 2012) and peaks were called using HOMER with an input ChIP-seq control. Overlapping FoxP3 ChIP-seq peaks were retained and ranked by signal intensity. The top 5000 ranked FoxP3 peaks were selected for a *de novo* motif analysis using MEME (Bailey et al., 2015). FIMO (Grant et al., 2011) was performed to identify variations in IR-FKHM motifs based on gap size and enrichment determined for non-consensus sequences from the 548 FoxP3 ChIP-seq sequences with an observed FKHM.

